# Modulation of nuclear and cytoplasmic mRNA fluctuations by time-dependent stimuli: analytical distributions

**DOI:** 10.1101/2022.02.25.481898

**Authors:** Tatiana Filatova, Nikola Popović, Ramon Grima

## Abstract

Temporal variation of environmental stimuli leads to changes in gene expression. Since the latter is noisy and since many reaction events occur between the birth and death of an mRNA molecule, it is of interest to understand how a stimulus affects the transcript numbers measured at various sub-cellular locations. Here, we construct a stochastic model describing the dynamics of signal-dependent gene expression and its propagation downstream of transcription. For any time-dependent stimulus and assuming bursty gene expression, we devise a procedure which allows us to obtain time-dependent distributions of mRNA numbers at various stages of its life-cycle, e.g. in its nascent form at the transcription site, post-splicing in the nucleus, and after it is exported to the cytoplasm. We also derive an expression for the error in the approximation whose accuracy is verified via stochastic simulation. We find that, depending on the frequency of oscillation and the time of measurement, a stimulus can lead to cytoplasmic amplification or attenuation of transcriptional noise.

## 1 Introduction

Many genes are transcribed in a bursty fashion [1], which is due to the fact that they spend most of their time in the “off” state, switching on for a relatively short time period during which a burst of mRNAs are rapidly produced. Furthermore, both the size of the burst in transcript numbers and the time between successive bursts are random [2, 3]. Noise can be due to intrinsic and extrinsic sources. Intrinsic noise stems from uncertainty in the timing of individual reaction events leading to transcription, whereas extrinsic noise arises independently of the gene but acts on it, e.g. through the number of RNA polymerases [4]. The mechanisms shaping transcriptional bursting are still not clearly understood and represent a topic of active research [5, 6].

The above has inspired the construction of stochastic models of gene expression with the aim of understanding how the distribution of mRNA numbers varies with transcriptional parameters. The simplest model that is in widespread use is the two-state telegraph model; considering exclusively intrinsic noise, its stochastic dynamics are described by the chemical master equation (CME) [7] which can be solved exactly [8, 9], yielding an explicit analytical solution for the distribution of transcript numbers as a function of the initiation rate, the switching rates between the “on” and “off” states of the gene, and the mRNA degradation rate. Within that model, the burst frequency is the rate at which the gene switches on, while the burst size is the initiation rate divided by the rate of switching off. Modifications of the telegraph model have been proposed to take into account noise in transcript numbers due to a wide variety of biological processes, such as the doubling of the gene copy number during DNA replication, partitioning of molecules between daughter cells during cell division, variability in the cell cycle duration time, coupling of gene expression to cell size or cell cycle phase, multiple off states, proximal-promoter pausing, RNA polymerase fluctuations, export from the nucleus to the cytoplasm, post-transcriptional modifications, and cell-to-cell variation in transcriptional parameters [10–21].

A common property of the bulk of published, analytically solvable models is their lack of description of the coupling of gene expression to an extracellular time-dependent signal. It is known that the identities and intensities of different stresses are transmitted by modulation of certain transcription factors (TFs) in the cytoplasm which exert an influence on gene expression upon their translocation to the nucleus [22–26]. This modulation is of particular importance in developmental biology whereby spatio-temporally varying distributions of TFs (morphogens) play a key role in establishing the body plan [27–32]. While TFs can exert influence on gene expression via modulation of the burst size and the burst frequency [33], modelling has shown that regulation through the latter is advantageous because weak TF binding is sufficient to elicit strong transcriptional responses [34]. In fact, the changes in distributions of nascent mRNA with TF concentration are very well captured by a telegraph model, modified so that the switching from the off state to the on state is an increasing function of the concentration [31, 35].

There are very few studies which have attempted to analytically solve the telegraph model (or similar models) for the distribution of transcript numbers in response to a time-dependent stimulus. In [36], an exact solution of the telegraph model with signal-dependent initiation rate is presented; because the initiation rate controls the burst size and not the frequency, that model does not capture the commonest way by which stimuli affect gene expression. In [37], an approximate solution of an auto-regulatory genetic feedback loop with signal-dependent initiation rate is presented which has the same disadvantage as mentioned for the previous study. By contrast, in [38], the stimulus is assumed to affect any one of the parameters in the telegraph model, which is hence compatible with the notion that stimuli are transmitted principally via modulation of the burst frequency; the solution of the resulting stochastic model is approximate and most accurate when the stimuli are slowly varying. In [39], a model where the burst frequency is modulated by an external signal is studied using a continuum approximation of the master equation. However, in these studies, there is no description of how the signal affects mRNA at different stages in its life-cycle, e.g. through differences between the temporal variation of the transcript numbers at the transcription site, in the nucleus, and the cytoplasm. That is important, as there are significant measured differences in the distributions and moments of nuclear and cytoplasmic mRNA [40, 41].

In this paper, we consider a stochastic model of gene expression in which a deterministic temporally-varying TF abundance modulates the burst frequency of a gene. By means of a novel approximation, we obtain closed-form analytical expressions for the time-dependent distribution of mRNA transcript numbers at any stage in the life-cycle, which is often correlated with a sub-cellular localization. The paper is organized as follows. In Section 2, we introduce a model of signal-dependent bursty gene expression where changes in some extracellular signal are reflected in the rate at which a gene switches on; for simplicity, we choose this rate to vary sinusoidally in time – an assumption that we relax later on. As the resulting model is multi-variable, the analytical time-dependent solution of its CME is highly challenging. The high dimensionality of the model here stems from the presence of *N* mRNA species, each describing mRNA abundance at a different stage in the life-cycle. We circumvent this challenge by postulating that the marginal time-dependent solution of mRNA at a particular life-cycle stage is described by an analytically tractable effective one-variable stochastic model with some unknown effective parameters. In Section 3, we describe a procedure by which the latter parameters can be found as a function of the parameters of the larger multi-variable model. We then show that for a wide range of parameters, the analytical solution of the one-variable model provides an excellent approximation to the marginal time-dependent distributions in the multi-variable model, as obtained via stochastic simulation. Finally, in Section 4, we extend the above procedure to models where the switching-on rate varies in a complex non-sinusoidal manner with time, which reflects the complexity of *in vivo* extracellular stimuli and the intricate molecular details of TF binding. We conclude with a discussion in Section 5.

## 2 Model description

In this section, we introduce a number of stochastic gene expression models of the mRNA life-cycle, which incorporate a temporally variable TF abundance due to an extracellular stimulus; see Fig. 1 for an illustration.

**Figure 1:**
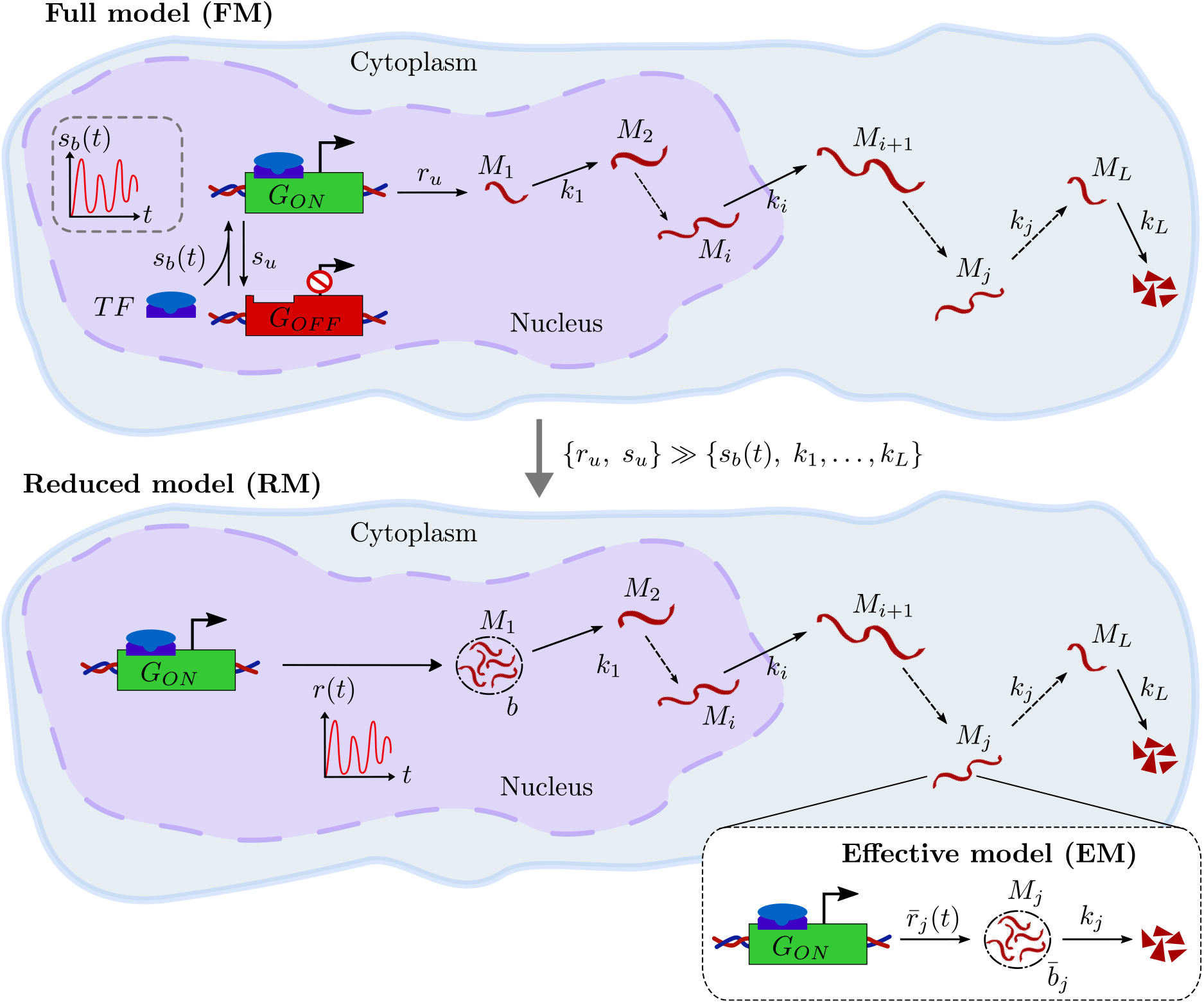
Illustration of three different stochastic models of the mRNA life-cycle. In the full model (FM), the gene can be in two states, active (*G*_*ON*_) and inactive (*G*_*OFF*_). Binding of TFs causes a transition from the inactive to the active state. The binding rate, *s*_*b*_(*t*), is time-dependent due to the temporal variation in TF numbers. The gene can switch back to its inactive state with rate *s*_*u*_. While the gene is active, transcriptional initiation occurs with constant rate *r*_*u*_, leading to synthesis of mRNA (*M*_1_). After the produced mRNA undergoes *L* stages of its life-cycle (*M*_*j*_) with rates *k*_*j*_ (*j* = 1, …, *L* − 1), it finally decays with rate *k*_*L*_. When transcription is bursty, the FM is well approximated by the reduced model (RM) which assumes that transcriptional initiation and gene inactivation rates (*r*_*u*_, *s*_*u*_) are much larger than the remaining kinetic rates. Here, mRNA synthesis occurs at a rate *r*(*t*) = *s*_*b*_(*t*) in bursts with mean size *b* = *r*_*u*_*/s*_*u*_. As before, the produced mRNA undergoes *L* stages of its life-cycle and eventually decays. In this paper, we show that the distribution of mRNA numbers in each life-cycle stage in the RM is well approximated by the distribution in an effective model (EM), which incorporates two rates: time-dependent bursty production of mRNA and degradation thereof. The advantage of the EM over the other two models is that it can be solved analytically, yielding time-dependent distributions of the mRNA in each life-cycle stage. See the main text for a more detailed description of these models.

### Full model (FM)

This model assumes that the gene can be either inactive (off), *G*_*OFF*_ or active (on), *G*_*ON*_, and that the mRNA life-cycle is divided into *L* stages, where the species *M*_*j*_ (*j* = 1, 2, …, *L*) denotes the mRNA in its *j*-th life-cycle stage. We assume that TFs can bind to enhancer or promoter sequences with some rate *σ*′ which leads to gene activation, at which point transcription is initiated. Furthermore, we assume that TF numbers vary periodically as *A*(1 +*ε* cos(*ωt*+*φ*)) where *A* is the amplitude, *ω* ≥ 0 denotes the frequency, and *φ* ∈ [− *π, π*) denotes the phase. Note that we choose the constant |*ε*| ≤ 1 such that the TF signal is always a positive-valued function. Note also that the choice of the cosine over a sine does not affect the periodicity of the signal, since cos(*x*) = sin(*x* + *π/*2) for all *x* ∈ ℝ. It then follows by the law of mass action that the activation rate from the inactive to the active state is given by

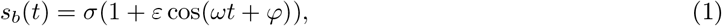

where *σ* = *Aσ*′. Note that we have assumed the binding rate to be a linear function of TF numbers here, which is a simplification, as TF binding kinetics is often cooperative [31], a case that we will discuss later on. Note also that, in reality, TF signals are to some degree noisy; however, in our case, we consider the signal to be deterministic for simplicity. The activation of the gene is a reversible reaction, i.e. the active gene can switch back to the inactive state with rate *s*_*u*_. Once the gene is activated, transcription initiation starts and with rate *r*_*u*_ leads to mRNA in stage one, denoted as *M*_1_. Subsequently, the synthesized mRNA progresses through its life-cycle by changing from state *M*_*j*_ to state *M*_*j*+1_ (*j* = 1, …, *L* − 1) with hopping rate, *k*_*j*_. At the end of its life-cycle, the final mRNA state, *M*_*L*_ decays with rate, *k*_*L*_. The system of chemical reactions describing the full model (FM) is given by

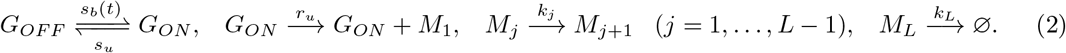

(Here, the empty set Ø denotes a sink of molecules.) Note that the mRNA life-cycle stages can represent any of the following processes: transcription initiation, splicing, elongation, maturation, and degradation [42]. As each of these can be modeled as one-step or multi-step processes, the parameter *L* is user-defined; hence, we present our analysis for the case of general *L* here. Note that if we define stages 1 to *R* to be nuclear, where *R* is some integer less than *L*, it follows that the time between initiation and export to the cytoplasm is a sum of *R* exponential random variables, each with mean 1*/k*_*j*_. Hence, the distribution of the nuclear retention time is a hypoexponential distribution; similarly, one can argue that the same distribution describes the lifetime of the cytoplasmic mRNA. Two special cases of this model have been previously studied: (i) the case *L* = 1 with pulse-like (non-sinusoidal) activation rate *s*_*b*_(*t*) [43], and (ii) the case of constant (non-time dependent) activation rate for general *L* [19].

### Reduced model (RM)

The analytical derivation of the time-dependent mRNA number distribution in a given stage *j* in the FM is a challenging task. A simplification is achieved from the observation that mRNA expression is often bursty, i.e. that the gene spends most of its time in the off state, producing a short-lived burst of molecules while in the on state [1, 3]. That observation leads us to introduce an simpler version of the full model, which we refer to as the reduced model (RM) and which is described by the reaction scheme

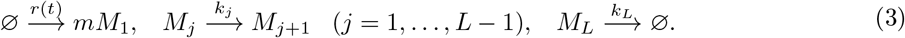

(Here, the empty set Ø denotes sources and sinks of molecules.) While there is no explicit gene switching in the RM, it is effectively taken into account by mRNA production occurring in bursts of size *m*, where *m* = 0, 1, … is a random variable chosen from a geometric distribution:

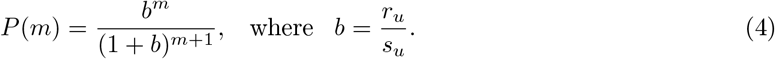

Note that the geometric distribution has a solid experimental and theoretical basis in the context of bursty expression [11, 44–48]. The parameter *b* is the mean burst size of mRNA molecules produced while the gene is active; values of this parameter for different genes have been reported in various studies [1]. In order to incorporate the dependence of transcription on TF numbers, we assume that burst production occurs with a time-dependent rate, which is exactly the gene activation rate from our FM, i.e. *r*(*t*) = *s*_*b*_(*t*).

As for the full model, *M*_1_ undergoes *L* life-cycle stages after it has been synthesized, followed by final mRNA degradation.

In Appendix B, we prove the equivalence of the two models when the transcriptional initiation rate and the gene inactivation rate of the FM are much larger than the remaining kinetic rate constants, i.e. when {*r*_*u*_, *s*_*u*_} ≫ {*s*_*b*_ (*t*), *k*_1_, …, *k*_*L*_}; therefore, in the remainder of the paper, our mathematical analysis is solely based on the RM.

To complete the setup of our RM, we introduce the following definitions. We define the vector of molecule numbers 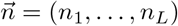, and we write ⟨*n*_*j*_⟩ (*j* = 1, …, *L*) for the average number of molecules of species *M*_*j*_. The RM can then be conveniently described by *L* species interacting via a set of *L* + 1 reactions with a rate function vector 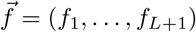 which has the following entries:

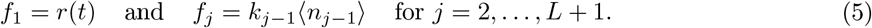

The rate functions *f*_*j*_ are the averaged propensities of the underlying chemical master equation (CME) [49]. The description of our model is completed by the *L* × (*L* + 1)-dimensional stoichiometric matrix **S**; the element *S*_*ij*_ of **S** gives the net change in the number of molecules of the *i*-th species when the *j*-th reaction occurs. Given the ordering of species and reactions as described in Eq. (3), it follows that the matrix **S** has the simple form

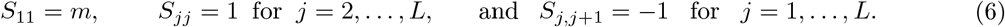

### Effective model (EM)

Finding the exact closed-form time-dependent mRNA distributions for each life-cycle stage in the RM is still very difficult. In this paper, we will show that we can well approximate the distribution of mRNA species in the *j*-th life-cycle stage in the RM – and, hence, in the FM – by the distribution in a simpler effective model (EM). The latter is defined by the following reaction scheme,

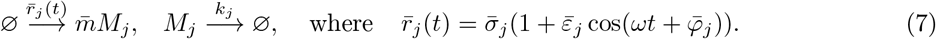

In the EM, 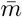 is a random variable chosen from the geometric distribution

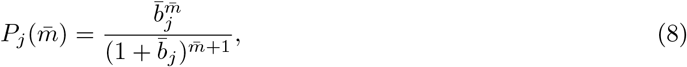

where 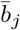 is the mean burst size for this model. Note that the subscript *j* in the parameters 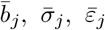, and 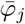 denotes their dependence on the life-cycle stage, *j* in the RM; these are to be determined later. However, we assume that the signal frequency, *ω* does not depend on the stage, *j*, and that it is the same as in the RM. Finally, the degradation rate of mRNA in the EM is *k*_*j*_, which is its hopping rate to the next stage in the RM – or its degradation rate if *j* = *L*.

## 3 Approximation of the distribution of mRNA numbers in the RM

Because of its high dimensionality due to the presence of *L* species, it is difficult to solve the CME for the RM and, hence, to obtain a time-dependent probability distribution of mRNA numbers in every stage in the life-cycle. We therefore take a different approach to obtain that distribution. In Section 3.1, we derive exact closed-form expressions for the first two moments of mRNA distributions in the RM. Subsequently, in Section 3.2, we find formulae for the effective parameters of the EM such that the mean number of mRNA in each life-cycle stage matches exactly that in the RM, while the variance in number fluctuations in the two models is matched approximately. Since the EM has the benefit that its CME can be solved exactly in time – as it has only one effective species – we finally obtain an analytical time-dependent distribution of mRNA numbers that is a good approximation of the distribution in the RM.

### 3.1 Exact closed-form expressions for mean and variance of mRNA distributions for the RM in the cyclo-stationary limit

In this section, we obtain analytical closed-form expressions for the first two moments of mRNA distributions in each life-cycle stage, in the limit of long times (*t* → ∞). Henceforth, we will refer to this limit as the “cyclo-stationary limit” [50], since for *t* → ∞, our solutions are still time-dependent functions due to the periodic time-dependent TF signal. Note that in our analysis we ignore fluctuations due to binomial partitioning at cell division; this approximation is valid as long as the mRNA lifetime is much shorter than the mean cell-cycle duration [18].

Given the CME describing the stochastic dynamics of the RM, it is straightforward to show from the corresponding moment equations [51] that the time evolution of the vector 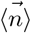 of mean molecule numbers is given by the set of ordinary differential equations (ODEs)

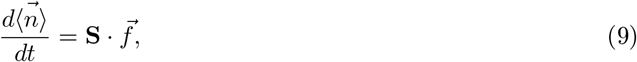

where 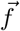 and **S** are defined in Eq. (5) and Eq. (6), respectively. Solving the above system of ODEs and taking the cyclo-stationary limit, we find that the solution can be written as

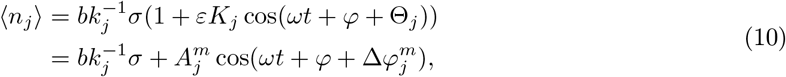

where 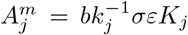 is the amplitude of the oscillation in the mean and 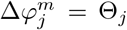 is the phase difference between this oscillation and the signal; the superscript *m* refers to the mean. The remaining parameters are defined as

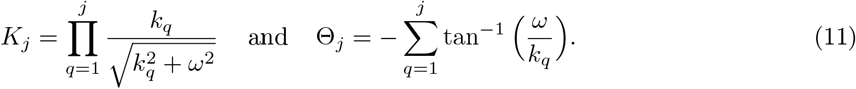

See Appendix C for a detailed derivation of these results. Since the propensities are linear in the number of molecules, the corresponding second moments at steady state are exactly given by a Lyapunov equation [51]. That equation, which is precisely the same as the one that is obtained from the linear noise approximation (LNA) [52], takes the form

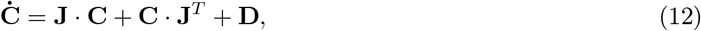

where the overdot denotes a time derivative. Here, **C, J**, and **D** are *L* × *L*-dimensional matrices: **C** is a covariance matrix that is symmetric (*C*_*ij*_ = *C*_*ji*_), **J** is the Jacobian matrix with elements 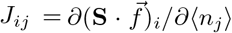, and 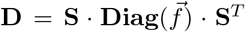 is a diffusion matrix, where 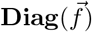 is a diagonal matrix whose elements are the entries in the rate function vector 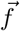. Eq. (12) can be solved explicitly for the covariance matrix **C**, the diagonal elements of which correspond to the variance of mRNA distributions in each life-cycle stage. In the cyclo-stationary limit, the latter are given by

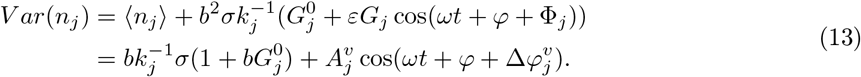

Here, 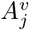 is the amplitude of oscillations in the variance, while 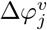 is the phase difference between these oscillations and the signal, which are obtained by solution of the equation

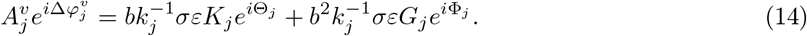

We also define 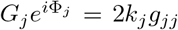 and 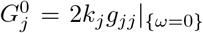, where the superscript 0 indicates that the expression is independent of the parameter *ω* and that it is real. Note that in the above expression, 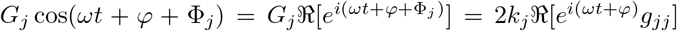, where ℜ[*z*] denotes the real part of the complex number, *z*. The function *g*_*ij*_ is given by the solution of the recurrence relation

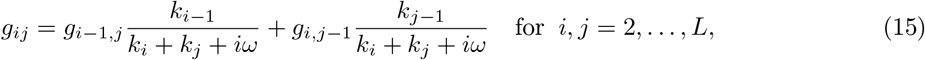

with initial conditions *g*_1*j*_ for *j* = 1, …, *L* defined as

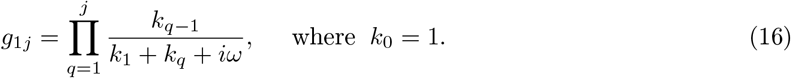

For detailed derivations of Eq. (13) and the solution of the recurrence relation in Eq. (15), we refer the reader to Appendix D. One can easily see that the mean stated in Eq. (10) and the variance given in Eq. (13) are periodic functions in time with the same period, *τ* = 2*π/ω* as the signal function.

In panels (a) and (b) of Fig. 2, we show the temporal evolution of the first two moments. In Fig. 2(c), we illustrate that the amplitudes of the moments are monotonically decreasing functions of the signal frequency, *ω*. Also, we show that, for some values of the frequency, the moment waves are in phase with the signal, as the phase differences become zero. While these results hold for every stage *j* in the mRNA life-cycle, we chose to present our plots in (c) for *j* = 14. In addition, it is clear that, while the amplitude of oscillations in the mean is lower than that of oscillations in the variance, the opposite is true for the phase difference of oscillations in the moments with respect to that of the signal, i.e. the variance wave always lags behind the mean wave which itself lags behind the signal. This interesting observation is clarified by a direct comparison of the two waves in Fig. 2(d).

**Figure 2:**
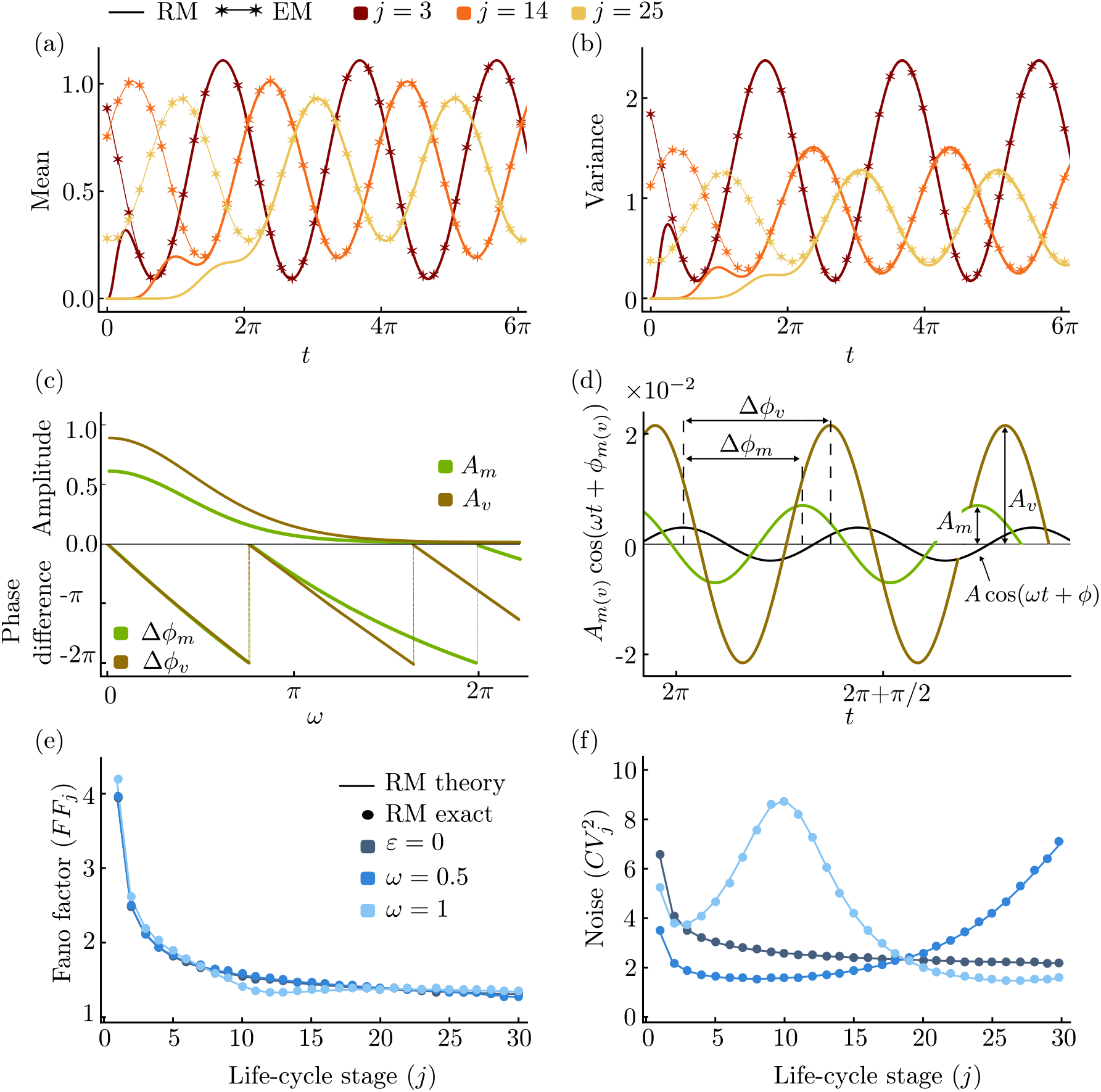
First two moments of the mRNA distributions in the RM. In panels (a) and (b), we illustrate the time evolution of the mean and the variance of mRNA distributions for three different stages of the mRNA life-cycle; thick solid lines show the direct numerical solution of the moment equations of the RM, given by Eq. (9) and Eq. (12), while thin solid lines with asterisks correspond to the approximation provided by the EM, Eq. (24). In panel (c), we show that the amplitudes of the oscillations in the moments decrease monotonically with signal frequency, *ω*; their phase differences with the signal reach zero for some value of *ω* at which the moment waves are in phase with the signal wave. In (d), we illustrate the time evolution of the time-dependent terms in the moments and the rescaled signal, 3 · 10^−3^ cos(*ωt* + *φ*), for *ω* = 3*π/*2. For panels (c) and (d), we used Eq. (10) and Eq. (13); we note that, while the decreasing behaviour of the amplitudes and phase differences holds for all stages *j* (*j* = 0, …, *L*), we chose to present it for *j* = 14. In panels (e) and (f), we show the variation of the Fano factor (*FF*_*j*_) and the noise 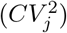 over the mRNA life-cycle for the RM. We present the case of constant signal, with *ε* = 0, and two examples of a time-dependent signal with different frequencies, as predicted by our theory (Eq. (18); solid lines) and simulation (SSA; dots). For the parameter values, see Appendix A.

One can easily show that if the TF signal frequency is much larger than the hopping rates, i.e. if *ω* ≫ *k*_*q*_ (*q* = 1, …, *j*), then the first two moments of the mRNA distributions are the same as in the case of a time-independent signal (*ε* = 0),

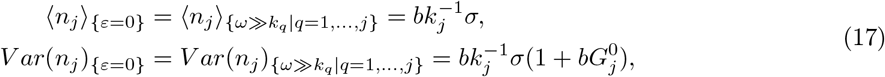

which is due to the fact that the amplitudes of the oscillations in the moments are decreasing functions of *ω*; see Fig. 2(c). Note that the time-averaged TF signal, 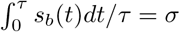, is the same as the TF signal when *ε* = 0. Hence, Eq. (17) implies that for high signal frequency, the mRNA only senses the constant time-averaged TF signal, which is in agreement with intuition.

We can also compute the Fano factor and the coefficient of variation squared. The first is a measure of how far distributions are from a Poissonian for which the Fano factor is 1, while the second is a measure of the magnitude of the noise. Analytical expressions for these quantities are obtained from the moment expressions in Eq. (10) and Eq. (13), and are given by

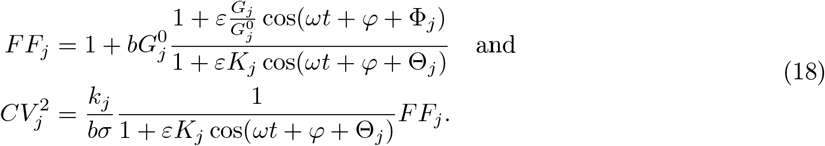

In Fig. 2(e), we show that for the case of identical hopping rates, with *k*_*j*_ independent of *j*, the Fano factor has an overall tendency to decrease as the mRNA progresses through its life-cycle, independent of the frequency and amplitude of the signal and of the measurement time, which is due to the factor in front of 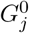 in Eq. (18) being approximately equal to one across parameter space: 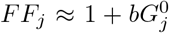. By contrast, in Fig. 2(f), we show that while the coefficient of variation squared is monotonically decreasing with life-cycle stage in the absence of an oscillatory signal, in its presence it can increase or decrease with the life-cell stage depending on the signal frequency and the time at which the measurement is taken. All results were verified using the stochastic simulation algorithm (SSA) for the RM – here, we applied a modified version of the Gillespie algorithm, which is described in Appendix E. If we associate the cytoplasm with stages *j* greater than some value, then this observation implies that a signal can either lead to cytoplasmic amplification of transcriptional noise (in the nucleus) or to its attenuation. While the results shown in panels (e) and (f) of Fig. 2 assume identical hopping rates, they also qualitatively hold when the hopping rates are non-identical.

### 3.2 Approximate mRNA distributions for the RM from the EM

While the moments to any order can be derived exactly for the RM, since all propensities are linear, the derivation of an expression for the marginal probability distribution of mRNA in each life-cycle stage proves to be a difficult challenge. Inspired by recent work which approximates the steady state solution of complex models of gene expression by that of simpler models [19], we seek to approximate the time-dependent solution of the multi-variable RM by the solution of the much simpler, one-variable EM. We proceed by finding the exact closed-form time-dependent solution for the mRNA distribution in the EM. In what follows, we consider that *j* has some fixed value between 1 and *L*, which is chosen by the user, according to which mRNA life-cycle stage one is interested in. Furthermore, we define *n* as the number of mRNA molecules of species *M*_*j*_ and *P*(*n*; *t*) as the probability of finding *n* molecules in the system at time, *t*. Given the reaction scheme of the EM from Eq. (7), it follows that the CME of our system is given by

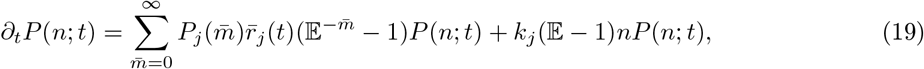

where 𝔼^*c*^[*P*(*n*)] = *P*(*n* + *c*), with *c* ∈ ℤ, denotes the standard step operator [7]. We define the generating function, 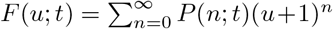 with *u* ∈ [−1, 0], to convert the above equation into the following partial differential equation (PDE):

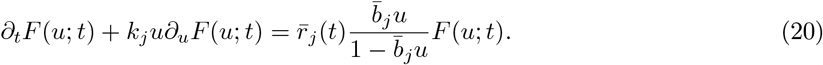

For *ω* ≠ 0, Eq. (20) admits the solution

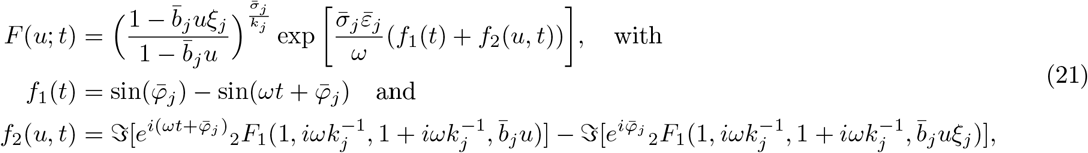

where _2_*F*_1_ is a hypergeometric function of the second kind [53, 54] and we have defined 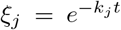; moreover, ℑ[*z*] denotes the imaginary part of a complex number, *z*. For the case of *ω* = 0, the solution of Eq. (20) is given by

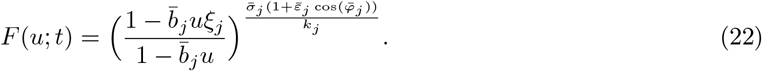

See Appendix F for a detailed derivation of these solutions. We note that the expressions in Eq. (21) and Eq. (22) are both well defined: first, 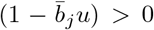 due to *u* ∈ [−1, 0], while 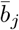 is some positive parameter; also, 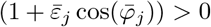 for 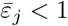, which follows from the definition in Eq. (26) below. The time-dependent marginal distribution is then found by using the formula

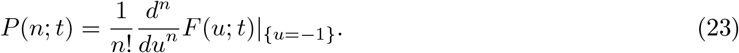

Here, we note that to obtain the solution in the cyclo-stationary limit, we just need to set *ξ*_*j*_ = 0 – in that limit, the solution (for *ω >* 0) will still be time-dependent. Note also that if the production rate is constant, i.e. if 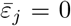 or *ω* = 0, then the solution reduces to a simple Negative Binomial (NB) distribution, which has been previously reported in [47, 55]. By contrast, when the production rate is time-dependent, the distribution of the mRNA species is not NB and can even be bimodal for some parameter values; see the supplementary Fig. F.1.

Having closed-form expressions for the mRNA distributions in the EM is not sufficient to approximate the mRNA distributions in the RM, since the parameters 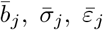, and 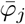 are unknown. We now seek to obtain analytical expressions for these unknown parameters by matching our expressions for the mean and the variance in the EM with the RM. From the solution for the generating function given in Eq. (21), it is straightforward to show that the first two moments of the EM in the cyclo-stationary limit can be written as

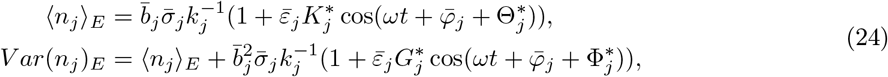

where the subscript *E* refers to the EM and the newly defined parameters are given by

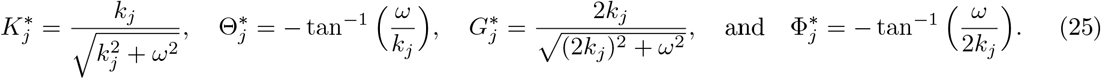

Our goal is to match the moments of mRNA distributions from the RM, as stated in Eq. (10) and Eq. (13), with the moments given in Eq. (24). Because of the complicated form of these analytical expressions, there might be more than one way of matching these moments; here, we present the most straightforward one that serves our purpose. First, we exactly match the means by setting ⟨*n*_*j*_⟩ = ⟨*n*_*j*_⟩_*E*_; by inspection, it is easy to verify that matching can be achieved by taking the constants in the EM to read

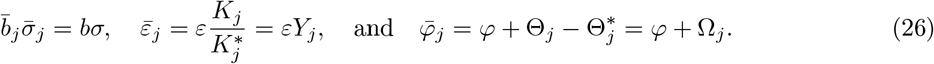

Given these parameters, it is not possible to match exactly the variances of the RM and the EM, except when *j* = 1 – the latter being obvious from an inspection of the reaction schemes of both models – or else when the TF signal is not time-dependent, i.e. when *ε* = 0 or *ω* = 0. (That case was studied in [19].) Note that in the limit of very large frequency *ω* – taken to be much larger than the hopping rates – one can also exactly match the second moments in the two models, which is due to the mRNA distributions in the RM being a function of the time-averaged TF signal only in that limit, as noted already. For the general case where *j* > 1, *ε* > 0, and *ω* > 0, one can match the time-independent terms in the expressions for the variance in the RM and the EM, which leads to the following additional constraints:

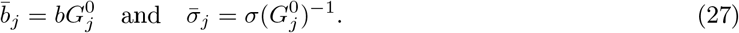

In summary, our approximate solution of the RM is given by Eq. (21), with the constants as defined in Eq. (26) and Eq. (27).

In panels (a) and (b) of Fig. 2, we compare the time evolution of the exact mean and variance in the RM (numerical solution of Eq. (9) and Eq. (12)) with the time evolution of the mean and variance, as computed from the EM model in the cyclo-stationary limit, Eq. (24). We observe excellent agreement between the two for various life-cycle stages. In Fig. 3, we verify that the distribution in the RM, as computed from stochastic simulation, is also in good agreement with the approximate distribution in the EM that is computed from the generating function given in Eq. (21), evaluated in the cyclo-stationary limit of *ξ*_*j*_ = 0.

**Figure 3:**
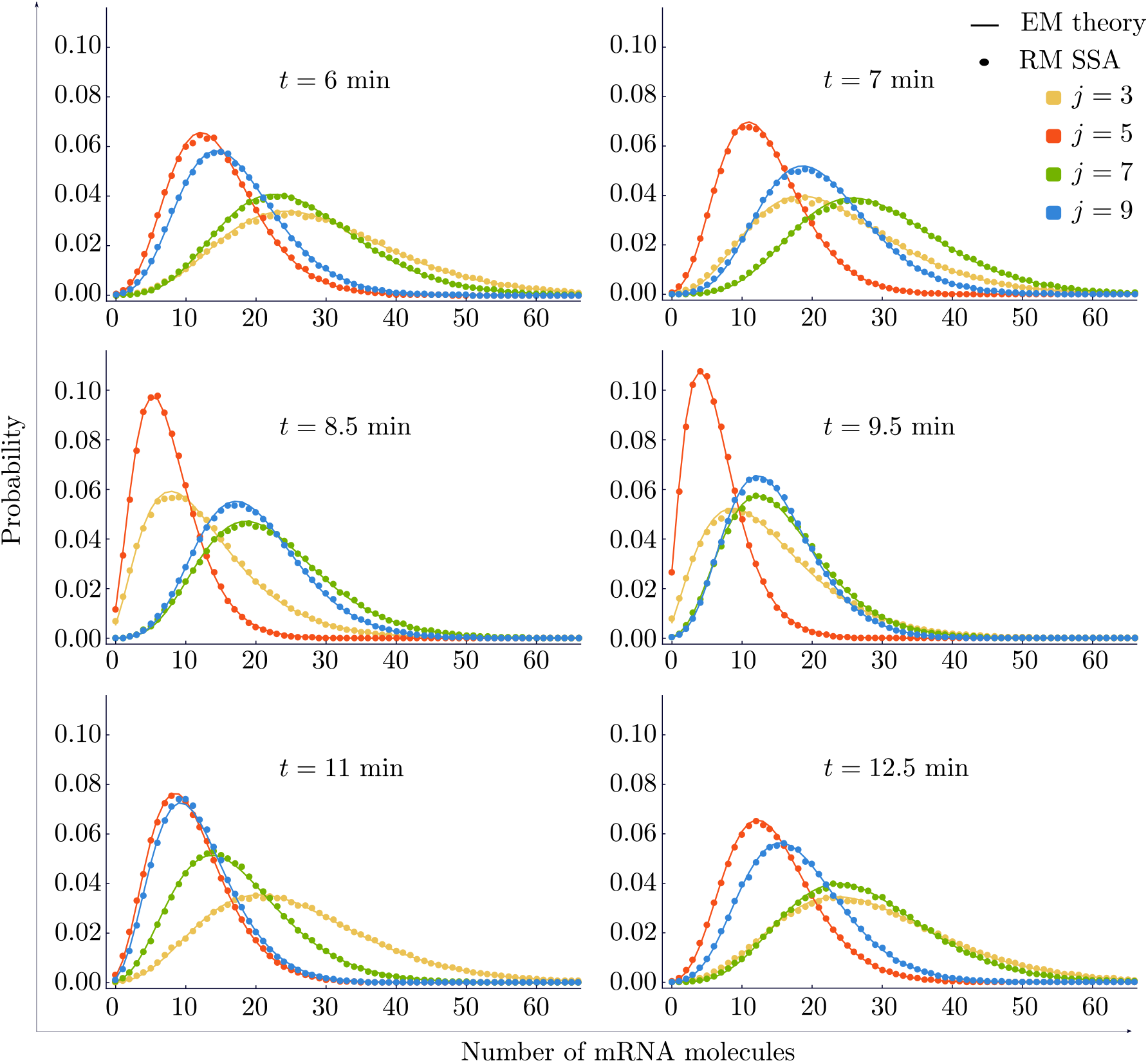
Comparison of the mRNA distributions in the RM with those in the EM. The mRNA distributions in the RM are not known analytically, and are hence computed from stochastic simulation (SSA; points). The mRNA distributions in the EM are given by Eq. (23), together with Eq. (21), evaluated in the cyclo-stationary limit and with the constants given in Eq. (26) and Eq. (27) (solid lines). We show the distributions for four different mRNA life-cycle stages (*j*) and six time points (*t*), as stated in the corresponding legends. These time points cover the time range of one signal period, *τ* = 2*π*. Note that for *j* = 1, our analytical distribution is exact (not shown), while for states with *j* > 1, the analytical distribution in the EM is a very good approximation to the distribution obtained from simulations of the RM. For the parameter values, see Appendix A.

### 3.3 Accuracy of the EM approximation

Next, we seek to investigate in detail the accuracy of the approximation to the RM that is provided by the EM. For each mRNA life-cycle stage, *j*, we define the vector *P* = (*p*_1_, …, *p*_*k*_) whose *i*-th entry, *p*_*i*_, is the probability of observing *i* mRNA molecules according to the EM; that probability can be determined from the generating function, Eq. (21). (Note that *k* is some integer which is assumed large enough such that *p*_*k*_ is very small.) Similarly, we define a vector *Q* = (*q*_1_, …, *q*_*k*_) whose *i*-th entry, *q*_*i*_, is the probability of observing *i* mRNA molecules according to the RM. The Hellinger distance (HD) between the probability distributions in the EM and the RM is then defined as

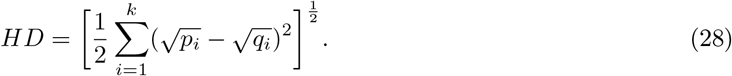

This measure of discrepancy between models, while ideal due to being based on the probability distributions, cannot be calculated analytically, since we do not have the exact analytical distribution for the RM. Hence, it can only be computed from stochastic simulation.

A different, analytical measure of the discrepancy between the RM and the EM is given by the relative error (*RE*) between the cyclo-stationary variance of the mRNA species predicted by both models. Using Eqs. (13) and (24), for each stage *j* we define the *RE* as

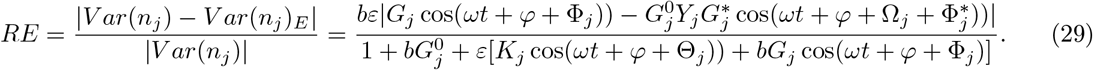

In Fig. 4, we investigate the relationship between the *HD* and the *RE* for different stages in the mRNA life-cycle across a wide range of parameter values. Three different points in parameter space, shown in panels (a) through (c) of Fig. 4, suggest that there is a linear relationship between the *HD* and the *RE*. This relationship between the two measures is confirmed in Fig. 4(d). Since we have an expression for the *RE*, it is hence easy to say when the EM provides a useful and accurate approximation of the distribution in the RM. We remark that the simple relationship between the *HD* and the *RE* is particular to the model under investigation, since generally one would expect the *HD* to depend on moments of order higher than two.

**Figure 4:**
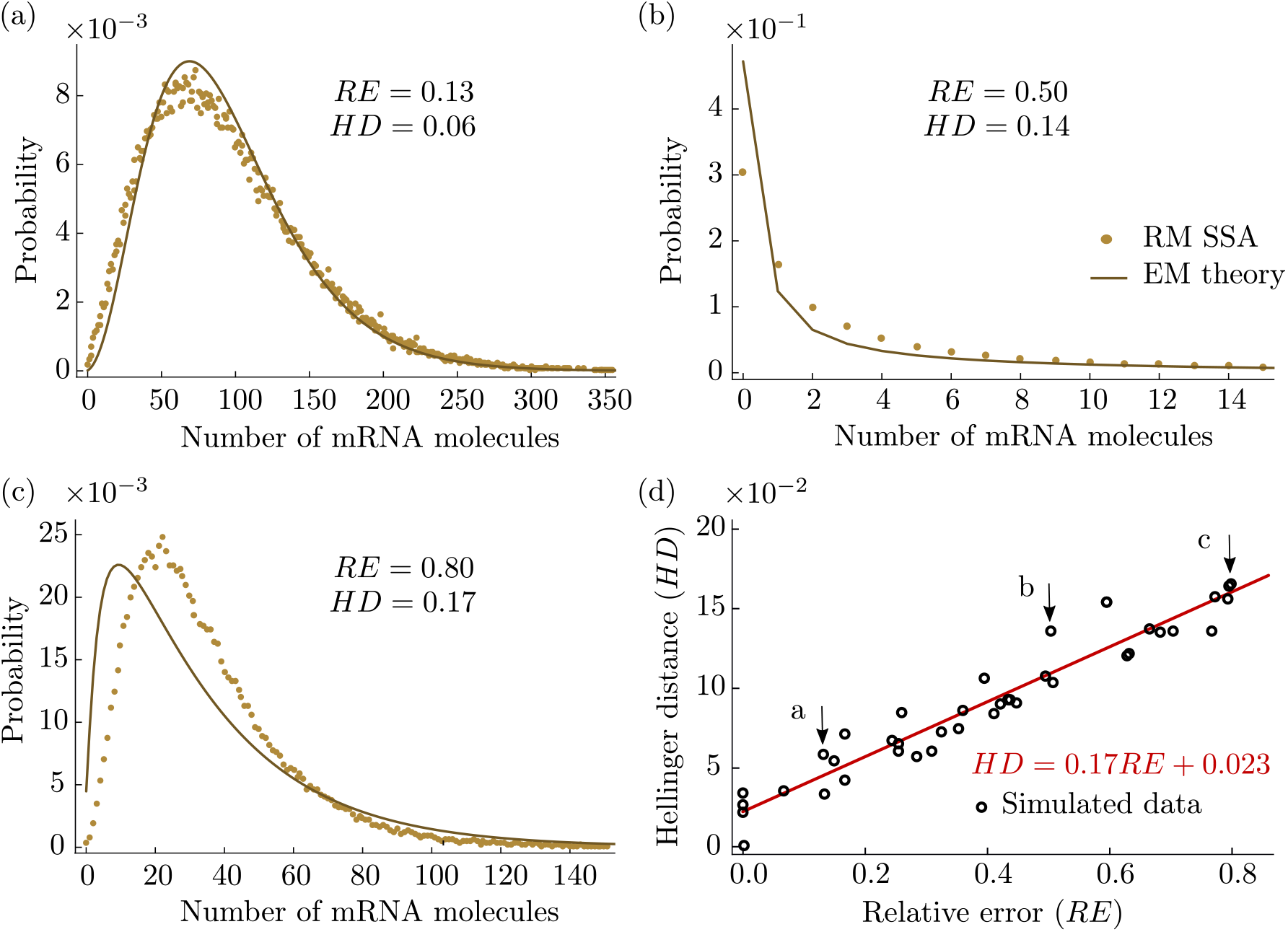
Accuracy of the EM approximation. In panels (a) through (c), we compare the distribution of mRNA numbers in the EM (Eq. (23) together with Eq. (21) and constants given in Eq. (26) and Eq. (27), computed in the cyclo-stationary limit; solid lines) and the RM (from stochastic simulation via the SSA; points). We also compute the *HD* between the two distributions and the *RE* of the variance of mRNA numbers in the EM. Comparison of (a) through (c) suggests that the *HD* increases linearly with the *RE*; this relationship is confirmed in panel (d) for 40 points and a fitted line *HD* = 0.17*RE* + 0.023 that is obtained by linear regression. We used Eq. (28) to calculate the *HD* and Eq. (29) to calculate the *RE*. For the parameter values, see Appendix A.

## 4 Generalization to the case of an arbitrary activation signal

Thus far, we have considered the approximation of the RM by the EM for the case when the activation rate from the inactive to the active state is given by *s*_*b*_(*t*) = *σ*(1 +*ε* cos(*ωt*+*φ*)). That approximation can be justified for the case of nuclear TF numbers varying in a sinusoidal manner assuming that the rate of switching is directly proportional to TF numbers, i.e. that there is no cooperativity. Of course, generally one expects the activation rate to have a much more complex time dependence, which is principally due to two factors: (i) environmental stimuli are coupled to gene expression via modulation of the number of TFs in the nucleus [24] – such stimuli will generally change in a complex time-varying manner, which will be reflected in TF numbers; (ii) binding of TFs to DNA is often cooperative [31], which implies that the rate of gene activation can be highly nonlinear in TF numbers via a Hill function dependence. To incorporate both of these factors, in this section we extend our results to the case of a general time-dependent activation rate that can be represented as a Fourier series:

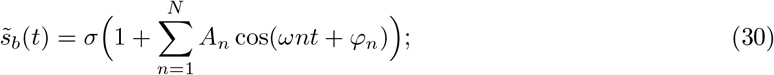

here, *N* ∈ ℕ_+_ and *A*_*n*_ ∈ ℝ for all *n* ∈ {1, …, *N*}, where *A*_*n*_ is such that 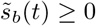. The RM is now given by Eq. (3) with 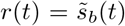. We hypothesize that the distribution of each state, *M*_*j*_, in the RM can be well approximated by a distribution in the EM defined in Eq. (7), where the production rate is now given by

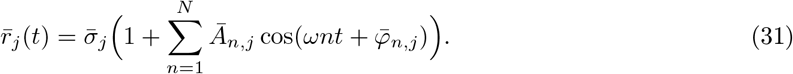

Note that the unknown parameters in this case are 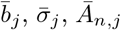 and 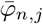, which we will determine below. All our derivations are performed in the cyclo-stationary limit of *t* → ∞. The exact closed-form solution of the probability generating function for the EM when *ω* ≠ 0 is given by

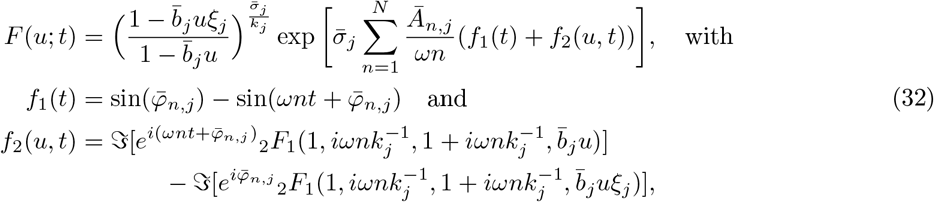

while for *ω* = 0, the solution reads

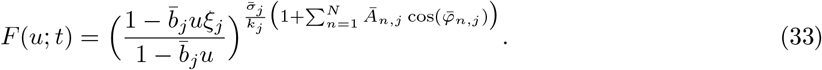

Here, _2_*F*_1_ is again the hypergeometric function of the second kind. The expressions in Eq. (32) and Eq. (33) are again well defined due to 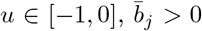, and 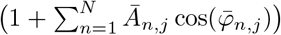 being positive for suitably chosen *Ā*_*n,j*_, which follows from the definition in Eq. (36) below. In order to derive analytical expressions for the unknown parameters, we follow the exact same steps in our mathematical analysis as in Section 3. First, we find the moments of the mRNA distributions in each life-cycle stage, *j*, for the RM and the EM. These are given by

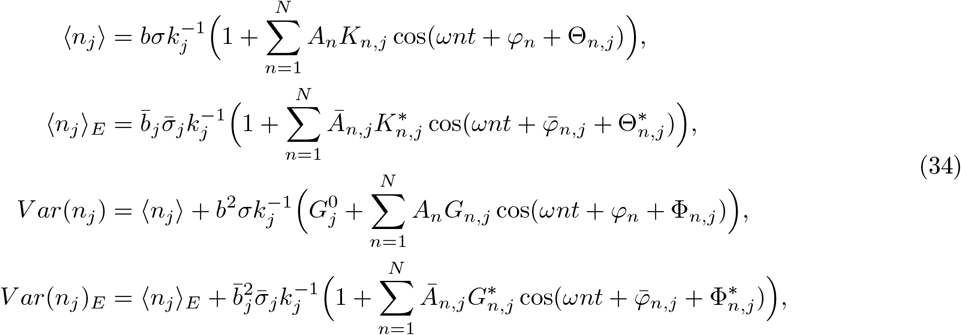

where the subscript *E* refers to the EM and the definition of the new parameters is as follows:

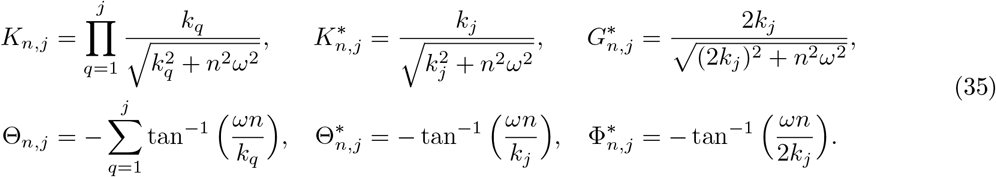

Also, we define 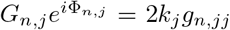 with *g*_*n,jj*_ = *g*_*jj*_|_*ω*↦*ωn*_, where *g*_*ij*_ is the solution of the recurrence relation in Eq. (15). By matching the moments in Eq. (34) in the same manner as in Section 3, we find the unknown parameters to be given by

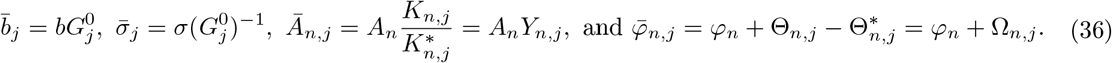

The approximate mRNA distribution in each mRNA life-cycle in the cyclo-stationary limit can be obtained by using 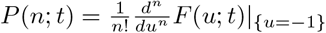 and Eq. (32), with parameters given as in Eq. (36).

In Fig. 5, we provide verification of the accuracy of the resulting generalized version of the EM by means of stochastic simulation. Here, we have used an approximately square wave for the time-dependent activation rate, as shown in Fig. 5(a), to model sharp TF pulses as considered in earlier work [43]. In panels (b) and (c) of Fig. 5, we show that the moments of mRNA distributions in the RM for four different stages in the mRNA life-cycle are well approximated by the moments of the EM. In panels (d) and (e) of Fig. 5, we verify that the approximate mRNA distributions in the EM are in excellent agreement with the mRNA distributions in the RM, which were computed using the SSA for two different time points.

**Figure 5:**
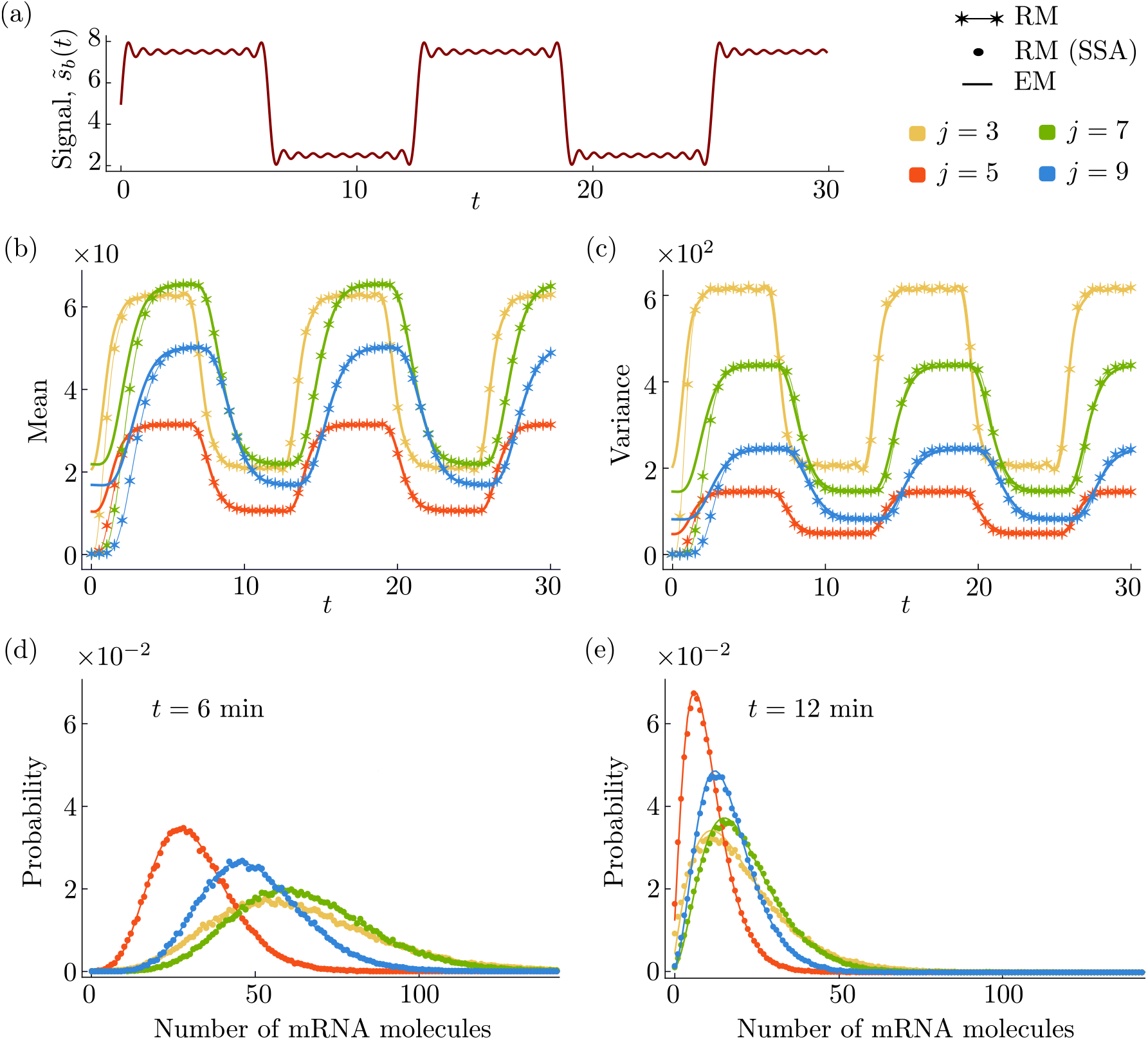
Generalization to the case of a general time-dependent activation signal. (a) A square wave-like time-dependent signal given by Eq. (30), where we specify *A*_*n*_ = −2*/*(*πn*) and *N* = 9. In panels (b) and (c), we compare the time evolution of the mean and the variance of the mRNA distributions in the RM (numerical solution of Eq. (9) and Eq. (12) with 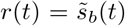 given by Eq. (30); thin lines with asterisks) with those in the EM as given by Eq. (34) (thick solid lines). In (d) and (e), we compare the mRNA distributions at time points *t* = 6 min and *t* = 12 min, respectively. The distributions are obtained from the SSA for the RM (points) and by Eq. (23) together with Eq. (32) and Eq. (36) for the EM (solid lines). The results in panels (b) through (e) are for four different stages in the mRNA life-cycle (*j*), as stated in the legends. For the parameter values, see Appendix A.

## 5 Conclusions

In this paper, we have considered a model for bursty transcription that is coupled to a time-varying extracellular stimulus, and we have applied a novel approximation to obtain the time-dependent distributions of mRNA in any life-cycle stage of interest. These stages often correspond to particular sub-cellular localization; hence, the model predicts, for example, the mRNA distribution at the transcription site, elsewhere in the nucleus, and in the cytoplasm – data that is accessible using experimental techniques [40,56]. We have shown that the resulting approximate distributions are in excellent agreement with stochastic simulation. In addition, we have found that the relative error between the true and the approximate variance of mRNA fluctuations – which can be calculated analytically – is directly proportional to the Hellinger distance between the distributions obtained from simulations and the theoretical ones. (The Hellinger distance is not accessible analytically, but only via simulation.) That relationship provides a convenient means to assess the accuracy of our theory without simulation.

Further, we have shown that apparent bimodality in the mRNA distributions can be generated by a time-varying stimulus when expression is bursty, which is interesting, considering that the solution of models of bursty expression without a stimulus gives a unimodal Negative Binomial distribution [11]. The intuition behind this phenomenon is clear, however: if the switching rate to the active transcription state is controlled by an oscillatory signal, then there are periods of intense transcription when the signal is very strong, whereas transcription almost switches off when the signal is weak. Our theory shows that if the stimulus is periodic, then (i) the oscillations in the variance of mRNA fluctuations lag behind those in the mean; (ii) the amplitude of oscillations in the first two moments decreases monotonically with the frequency; (ii) the Fano factor of mRNA fluctuations tends to decrease with life-cycle stage; (iii) the noise in mRNA fluctuations, as quantified by the coefficient of variation squared, can increase or decrease with life-cycle stage depending on the time of measurement and the frequency of the stimulus. The latter implies that the stimulus can either lead to an apparent increase of the noise in the cytoplasm compared to that in the nucleus, or to the opposite case of attenuation. We are not aware of experimental data that can verify these predictions, since observations reported in the literature were made in the absence of a time-varying stimulus [40, 41].

We note that other theoretical studies have sought to derive closed-form time-dependent mRNA distributions for various models of gene expression. These can be classified as follows: (i) those which do not consider a time-varying stimulus [11, 57–61], in that they study how gene expression approaches steady state given a perturbation that is applied at a point in time, e.g. with the initial condition given by mRNA numbers following cell division – in that case, the kinetic rates do not vary with time; (ii) those which consider a stimulus that varies with time [36–39]. The major difference between our paper and the latter is that we derive time-dependent analytical distributions for the mRNA at any stage of its life-cycle which often correspond to specific sub-cellular localization. For example, if we choose the simplest case of *L* = 2 in our model, then the distributions of *M*_1_ and *M*_2_ can be interpreted as being for nuclear and cytoplasmic mRNA, which would be under the assumption that the time from initiation to a mature mRNA appearing in the nucleus is exponentially distributed, as is the time for export from the nucleus to the cytoplasm. Deviations from the exponential assumption can also be easily incorporated into our framework. For example, if the distribution of the export time is Erlang with shape parameter *k*, then one could apply our model with *L* = *k* + 3, where *M*_1_ is nuclear mRNA, *M*_*k*+3_ is the cytoplasmic mRNA, and *M*_2_, …, *M*_*k*+2_ are dummy species introduced to capture the Erlang distributed delay. Another interesting application of our model would be to predict the distribution of bound RNA polymerase (RNAP) along the gene, in response to a time-varying stimulus; in that case, under the assumption that volume exclusion is not significant, the species *M*_*i*_ can be interpreted as the RNAP on gene segment *i* [16]. Besides the forward predictive power of our theory, the practical use thereof might lie in the resulting theoretical distributions, together with likelihood-based inference methods [62, 63], as a reliable means for estimating kinetic parameters from experimental population snapshot data of nascent, nuclear, and cytoplasmic mRNA measured for time-varying extracellular stimuli.

## Acknowledgments

T.F. acknowledges useful discussions with Juraj Szavits-Nossan. This work was supported by a departmental PhD studentship to T.F.

## Appendix

### A Parameter values and other details of the figures

Fig. 2 For all panels, we have arbitrarily chosen the number of mRNA stages to be *L* = 30. In panels (c) and (d), we have used the parameter *σ* = 1*/ε*, with *ε* = 0.9. In panels (e) and (f), we present our results for the time point *t* = 10 min, while in panel (f) we additionally include results for the time point *t* = 11.5 min. We only present the Fano factor for *t* = 10 min, since it is negligibly affected by time. We also note that for the case of *ε* = 0, which indicates a constant signal, both the Fano factor and the noise are independent of time. The parameter values that have been used in all panels are *σ* = 1 min^−1^, *ε* = 0.9, *b* = 3, *φ* = *π/*2, *ω* = 1 min^−1^, and *k*_*j*_ = 5 min^−1^ (*j* = 0, …, *L*), unless otherwise stated.

Fig. 3 The parameter values that have been used in all panels are *b* = 10, *ε* = 0.5, *σ* = 5 min^−1^, *ω* = 1 min^−1^, and *φ* = *π/*2, as well as *k*_1_ = 4.2, *k*_2_ = 2.6, *k*_3_ = 2.4, *k*_4_ = 3.7, *k*_5_ = 4.8, *k*_6_ = 2.7, *k*_7_ = 2.3, *k*_8_ = 3.7, and *k*_9_ = 3.0, all of which have units of min^−1^.

Fig. 4 The parameter values that have been used in panel (a) are *b* = 50, *ω* = 1 min^−1^, *ε* = 0.99, *t* = 9.2 min, and *j* = 2. The parameter values that have been used in panel (b) are *b* = 150, *ω* = 0.22 min^−1^, *ε* = 0.995, *t* = 9.9 min, and *j* = 8. The parameter values that have been used in panel (c) are *b* = 150, *ω* = 1 min^−1^, *ε* = 0.99, *t* = 9.2 min, and *j* = 5. In order to obtain the simulated data (points) in panel (d), we performed stochastic simulations over a range of parameter space (*b, ω, ε*) ∈ [1, 150] × [0, 5] × [0, 1], for *L* = 10 mRNA life-cycle stages and for two time points, *t* = 9.2 min and *t* = 9.9 min. Then, we randomly chose 40 points to present from the resulting data. The remaining parameters that have been used for panels (a) through (d) are the same, and are given by *σ* = 5 min^−1^ and *φ* = *π/*2, as well as by *k*_1_ = 4.2, *k*_2_ = 2.6, *k*_3_ = 2.4, *k*_4_ = 3.7, *k*_5_ = 4.8, *k*_6_ = 2.7, *k*_7_ = 2.3, *k*_8_ = 3.7, and *k*_9_ = 3.0, all of which have units of min^−1^.

Fig. 5 The parameter values in all panels are as in Fig. 3, with the exception of *b* = 20 and *ω* = 0.5 min^−1^.

### B Equivalence of the full model (FM) and the reduced model (RM) under timescale separation

In this section, we will show that in the limit of {*r*_*u*_, *s*_*u*_} ≫ {*s*_*b*_(*t*), *k*_1_, …, *k*_*L*_}, the mRNA distributions obtained from the RM are exactly the same as those in the FM. For the FM, we consider the system of chemical reactions given in Eq. (2). We define 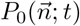 to be the probability of finding 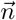 molecules in the system at time *t* when promoter is active and 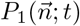 to be the probability when it is inactive. The vector of the number of mRNA molecules in each life-cycle stage is defined as 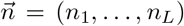. The CME is then given by the set of coupled equations

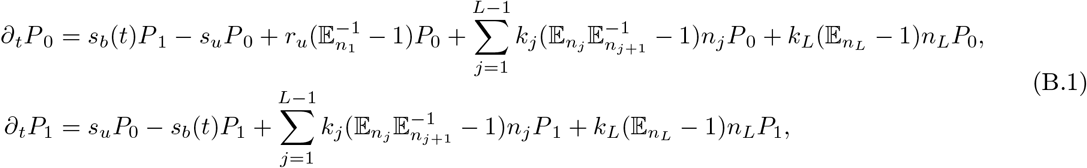

where 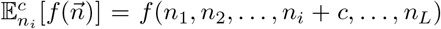, with *c* ∈ ℤ, denotes the standard step operator. Now, we define the corresponding probability-generating functions as

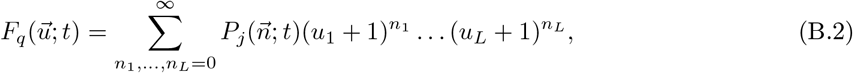

for *q* = 0, 1; here, 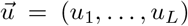 is a vector of real variables corresponding to the state 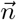, with *u*_*q*_ ∈ [−1, 0] for *q* = 1, …, *L*. Hence, we can rewrite the above CME as the system of PDEs

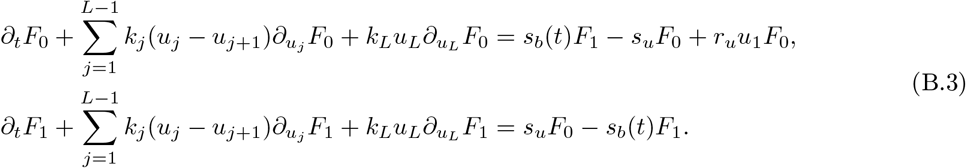

Next, using the method of characteristics, we convert the above PDEs into the following system of ordinary differential equations (ODEs):

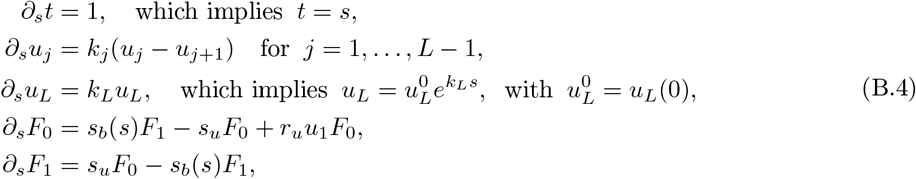

where *s* ∈ ℝ is the characteristic variable. Using *x*(*s*) = 1 + *ε* cos(*ωs* + *φ*), we rewrite the above ODEs for the generating functions as

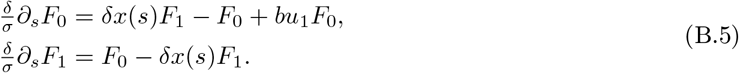

where *b* = *r*_*u*_*/s*_*u*_ and *δ* = *σ/s*_*u*_. We assume that *s*_*u*_ ≫ 2*σ*, which implies that the promoter spends most of its time in the inactive state, since *s*_*u*_ ≫ 2*σ* ≥ *s*_*b*_(*t*). It follows that *δ* ≪ 1 can be taken as a small perturbation parameter. Also, we assume that the parameter *r*_*u*_ is of the same order of magnitude as *s*_*u*_ such that *b* remains constant as *δ* becomes very small. Here, we note that by assuming *r, s*_*u*_ ≫ *s*_*b*_(*t*), we automatically also assume that the parameters *k*_*j*_ (*j* = 1, …, *L*) are of the same order of magnitude as the parameter *σ*: *r, s*_*u*_ ≫ *k*_*j*_. We may then take *F*_*q*_ (*q* = 0, 1) to have a series expansion in *δ*:

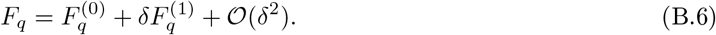

Substituting the above expansion into Eq. (B.5) and collecting leading-order terms in *δ*, i.e. terms of the order *δ*^0^, we obtain 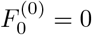. Similarly, collecting first-order terms in *δ*, we find the system

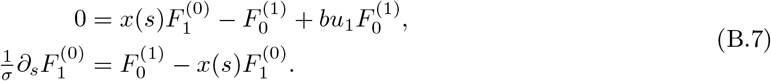

Using *F* = *F*_0_ + *F*_1_ and Eq. (B.7) we obtain the following ODE for *F*^(0)^:

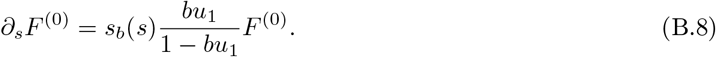

Eq. (B.8), together with Eq. (B.4), then gives us the following system:

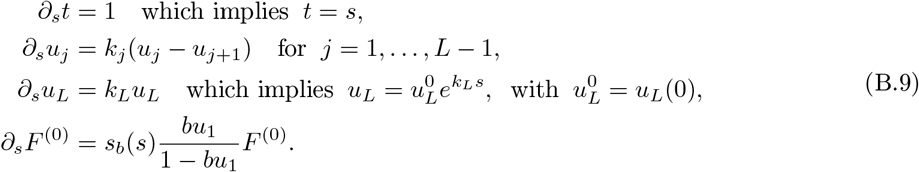

Now, for the RM, we consider the system of chemical reactions given in Eq. (3). We define 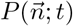 to be the corresponding new probability. Then, the CME for our system is given by

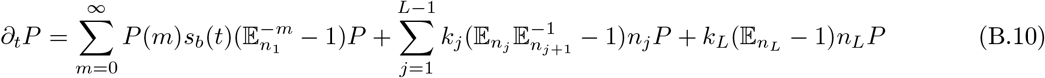

where *P*(*m*) = *b*^*m*^*/*(1 + *b*)^*m*+1^ (*m* = 0, 1, …) is a geometric distribution with mean, *b*. Then, by using the method of generating functions, we obtain the following PDE;

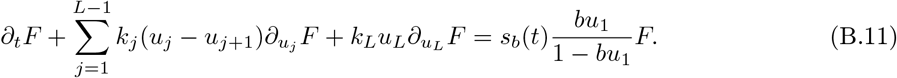

Since the solution for *F*^(0)^ from Eq. (B.9) and the solution for *F* from Eq. (B.11) are identical, we conclude that the mRNA distributions obtained from the FM and the RM are the same under the timescale separation assumed here.

### C Closed-form expressions for the mean number of mRNA molecules in each life-cycle stage in the RM

In this section, we present a detailed derivation of the solution to Eq. (9) in the cyclo-stationary limit. The governing system of differential equations can be written as

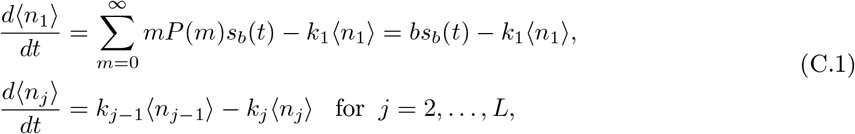

where we have used the definition *r*(*t*) = *s*_*b*_(*t*). We apply the Laplace transform to the above equations and obtain the system

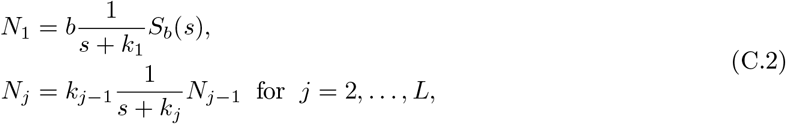

where 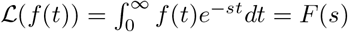 is the Laplace transform and ℒ(⟨*n*_*j*_⟩) = *N*_*j*_, as well as ℒ(*s*_*b*_(*t*)) = *S*_*b*_(*s*). Here, we have used the initial conditions ⟨*n*_*j*_⟩|_*t*=0_ = 0 for *j* = 1, …, *L*, which indicates zero mRNA molecules in the system initially. Now, we apply the inverse Laplace transform, ℒ^−1^(*F*(*s*)) = *f*(*t*), to Eq. (C.2) to obtain the following system:

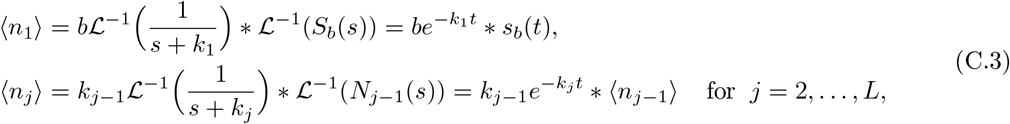

where ∗ denotes the convolution operator. Then, the system in Eq. (C.3) can be written as

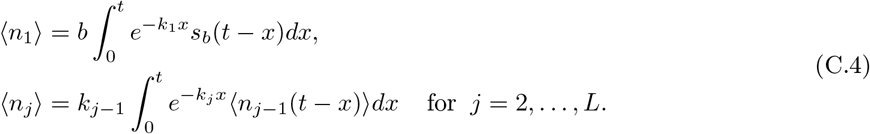

Evaluating the first integral in the cyclo-stationary limit of *t* → ∞, we obtain

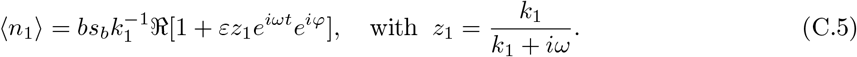

Note that we have expressed the time-dependent signal in the form *s*_*b*_(*t*) = *σ*ℜ[1 + *εe*^*iωt*^*e*^*iφ*^] here, where ℜ[*z*] again denotes the real part of the complex number, *z*. Similarly, we can find the solution for each ⟨*n*_*j*_⟩ from Eq. (C.4) in the limit of large times, where we also use the solution for ⟨*n*_1_⟩ from Eq. (C.5). Then, one can easily show that for *j* = 2, …, *L*,

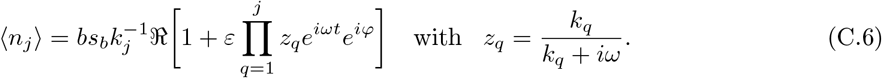

We can simplify the expressions in Eq. (C.5) and Eq. (C.6) by expressing the complex numbers *z*_*q*_ in polar form as

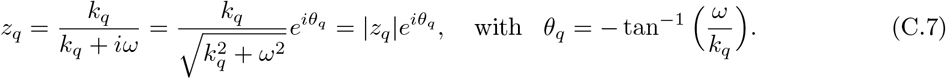

Then, we use the identity

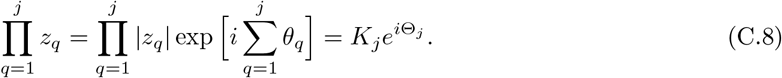

Hence, Eq. (C.5) and Eq. (C.6) simplify to

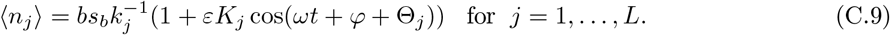

Since Eq. (C.9) represents a wave for each stage *j*, we can also rewrite the above as

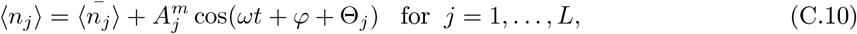

where 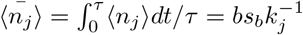 is the time-averaged mean over one period of time, 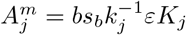 is the amplitude, and *φ* + Θ_*j*_ is the phase of the time-dependent oscillatory part.

Here, we note that one can easily show that the amplitude 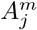 is a decreasing function of *ω*. Also, for *ω* ≫ *k*_*q*_ (*q* = 1, …, *j*), it follows that 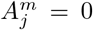, i.e. that the mean is constant and equal to the time-averaged mean. Additionally, we have that

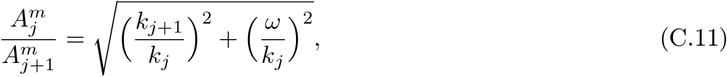

which indicates that the amplitude can increase or decrease with the life-cycle stage *j*, depending on the values of the parameters *k*_*j*+1_, *k*_*j*_, and *ω*.

### D Exact solution of the Lyapunov equation for the RM

The Lyapunov equation mentioned in Eq. (12) can be solved explicitly for the covariance matrix **C** in the cyclo-stationary limit. The non-zero elements of **J** are given by

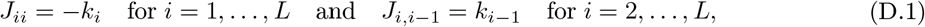

while the non-zero elements *D*_*ij*_ of **D** read

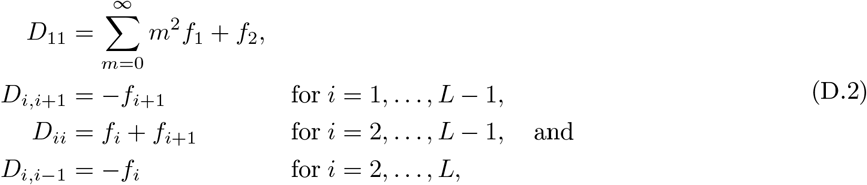

where *f*_*i*_ is defined in Eq. (5) and given in Eq. (C.9) when evaluated at ⟨*n*_*i*_⟩. Also, we have that

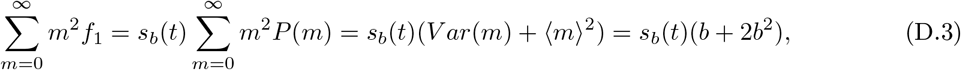

because *P*(*m*) is a geometric distribution with mean burst size *b*. With these definitions, we can express the Lyapunov equation as a system of *L*^2^ differential equations:

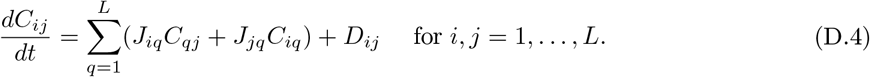

Considering only the non-zero elements of **J**, Eq. (D.4) simplifies to

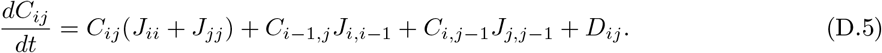

A further simplification is achieved by considering only the non-zero elements of **D**:

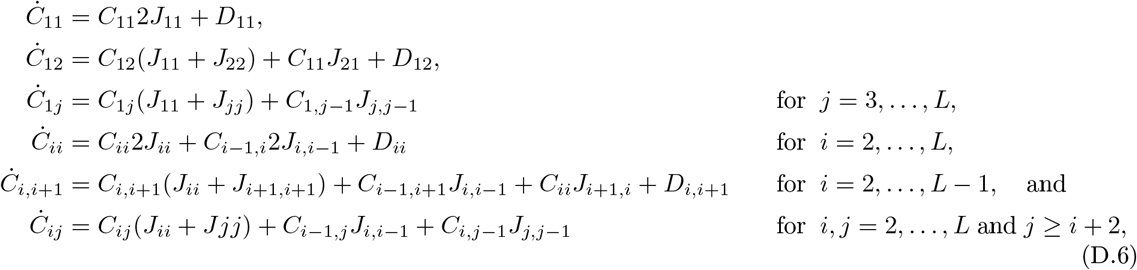

where the overdot denotes differentiation with respect to time *t*. Substituting the definitions of *J*_*ij*_ and *D*_*ij*_ into Eq. (D.6), we obtain

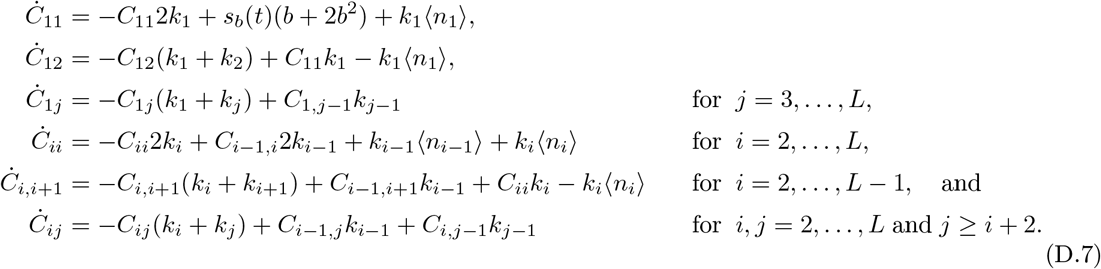

Now, we apply the Laplace transform to Eq. (D.7) with ℒ(⟨*n*_*j*_⟩) = *N*_*j*_, ℒ(*s*_*b*_(*t*)) = *S*_*b*_(*s*), and ℒ(*C*_*ij*_) = *c*_*ij*_, which gives us the following system of equations:

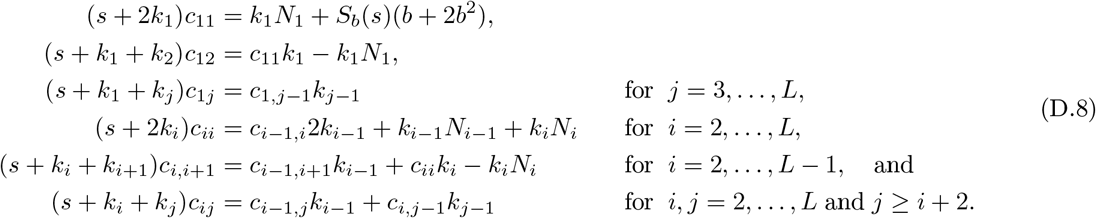

Here, we have used the initial conditions ⟨*n*_*j*_⟩|_*t*=0_ = 0 and *C*_*ij*_|_*t*=0_ = 0 for *i, j* = 1, …, *L*. Now, in Laplace space, we can use the expressions from Eq. (C.2), which gives us

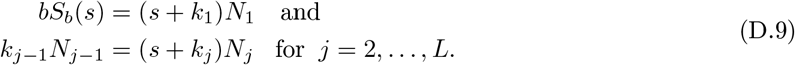

We substitute the above expressions into Eq. (D.8) and hence obtain the following simplified system:

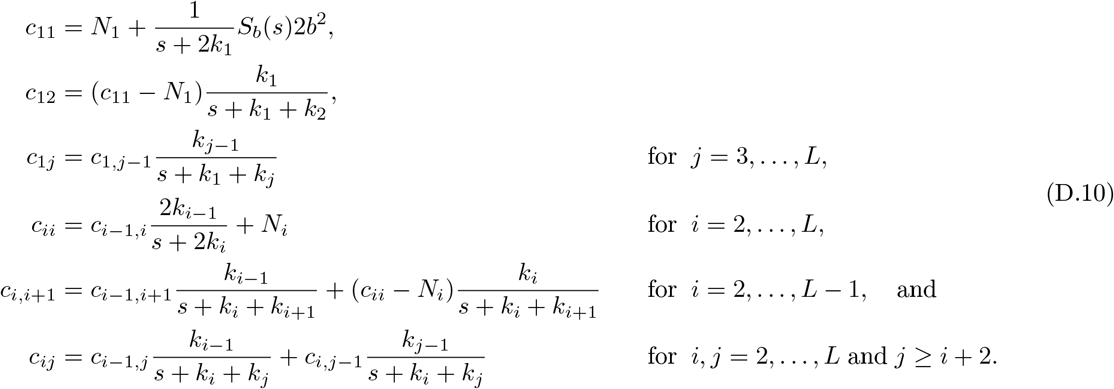

Now, we define new functions *f*_*ij*_ such that

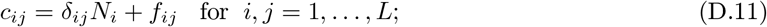

then, the system in Eq. (D.10) transforms to

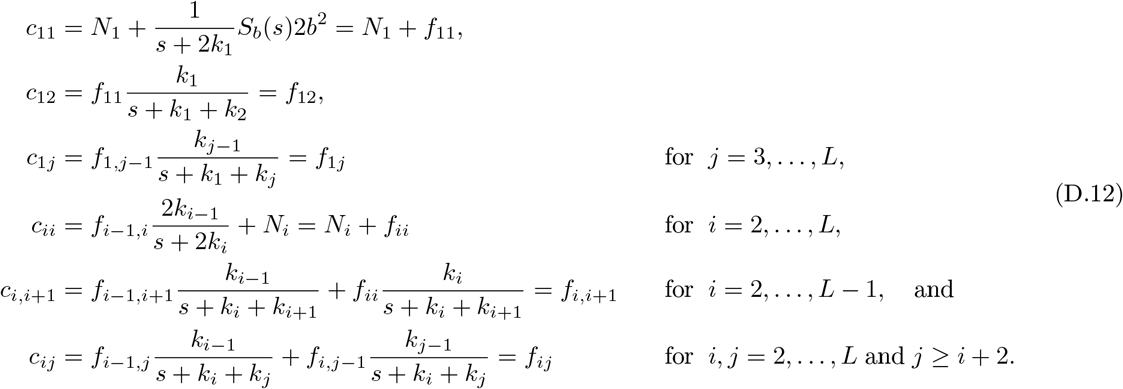

From the above transformation, it is clear that the function *f*_*ij*_ is naturally defined as

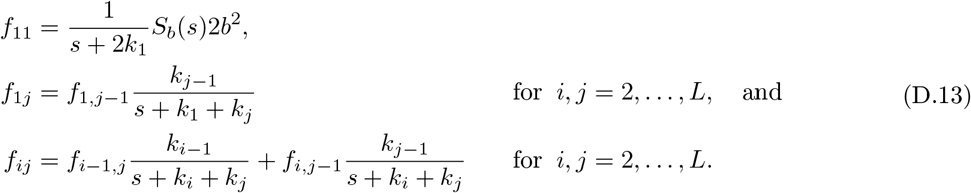

Next, we apply the inverse Laplace transform to Eq. (D.11) and obtain the expression

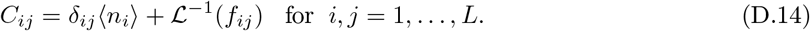

For the full solution of the covariance matrix **C**, we need to find the expressions for *F*_*ij*_ =ℒ^−1^(*f*_*ij*_). By going through the same steps as in Section C – use of the convolution property and introduction of a time-dependent signal in the form *s*_*b*_(*t*) = *σ*ℜ[1 + *εe*^*iωt*^*e*^*iφ*^], followed by evaluation of the integral over [0, ∞) to enforce the cyclo-stationary limit – we find that

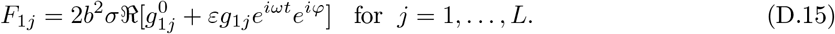

Here, *g*_1*j*_ is a complex function that is defined as

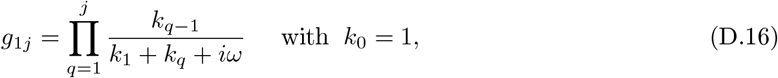

and 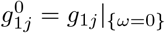. (The superscript 0 indicates that the expression is independent of the parameter *ω* and that it is a real function.) Now, we assume that

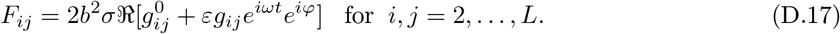

By combining the third equation in Eq. (D.13) and Eq. (D.17), one can easily find the recurrence relation

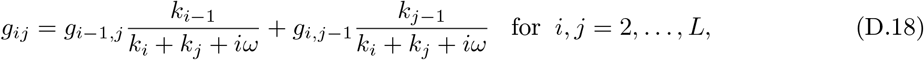

with initial conditions *g*_1*j*_ as defined in Eq. (D.16). Summarising our results, we have that the elements of the covariance matrix **C** for *i, j* = 1, …, *L* have the following closed-form expressions:

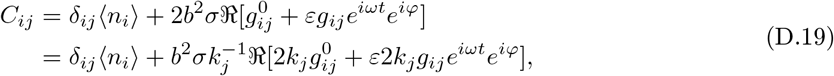

where the complex function *g*_*ij*_ is defined by the recurrence relation in Eq. (D.18) and 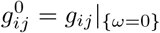 is a real function. The solution of this recurrence relation is given by

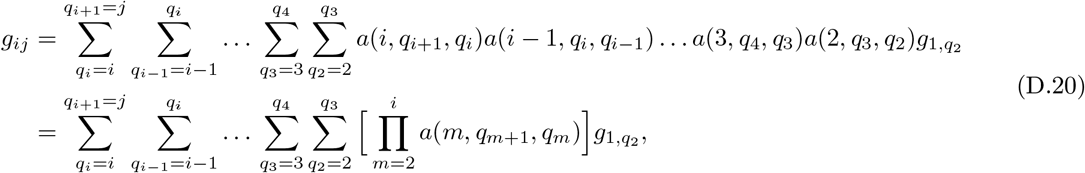

where *m* = 2, 3, …, *i, q*_*m*+1_ ≥ *m, m* ≤ *q*_*m*_ ≤ *q*_*m*+1_, and

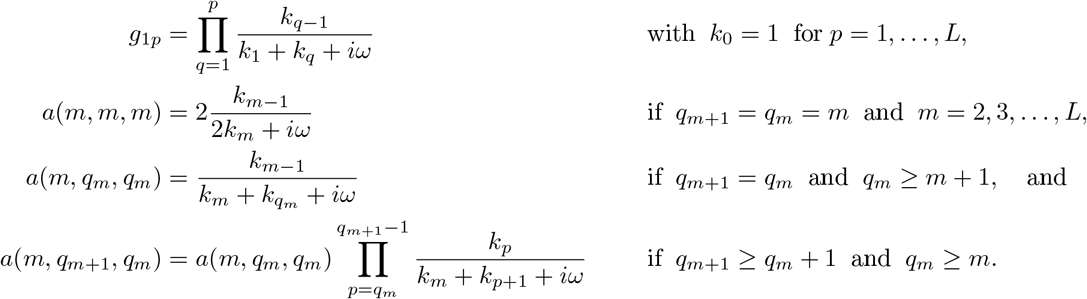

For *i* = *j*, we can obtain expressions for the variance from Eq. (D.19), which are given by

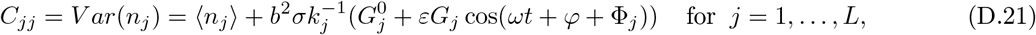

where 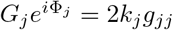 and 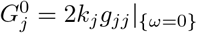.

### E The modified stochastic simulation algorithm (SSA)

For the stochastic simulations of our model as presented in the paper, we used the modified Gillespie algorithm for time-dependent propensity functions [64]. The main steps in the algorithm for generating one trajectory are as follows:

1. Initialize the time *t* = 0 and the number of molecules of each species, 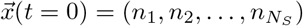, where *N*_*S*_ is the total number of species, and perform the following steps as long as *t* ≤ *t*_max_.
2. Generate two independent uniform random numbers on the interval (0, 1): *r*_1_, which defines the time that passes until the next reaction occurs; and *r*_2_, which defines the next reaction that occurs.
3. The time that passes until the next reaction occurs, Δ, is exponentially distributed with parameter

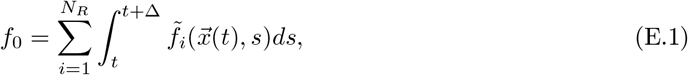

where the propensities 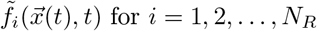 for *i* = 1, 2, …, *N*_*R*_ depend explicitly on time and *N*_*R*_ is the total number of reactions. Note that 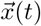 is constant in the above integral, as no reactions take place within the time interval [*t, t* + Δ). In our case, the above integral can be solved analytically; we write the propensity functions of the RM as

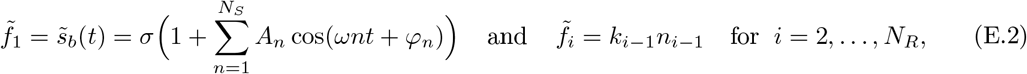

where we have *N*_*R*_ = *L* + 1 for our model, with *n*_*i*−1_ the entries of the array 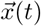. Here, we have considered the general case for our time-dependent signal, which can been written in Fourier form; see Section 4, with 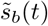 as in Eq. (30). Note that in our case, only the propensity function 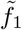 is explicitly dependent on time. The solution of the integral in Eq. (E.1) is then given by

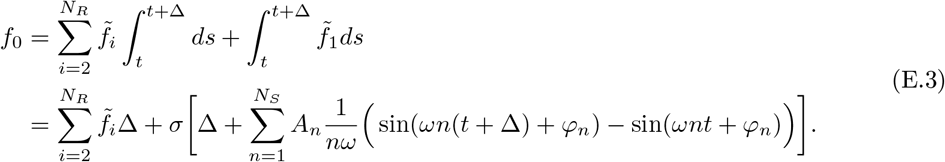

In order to obtain the time interval Δ, we solve the algebraic equation

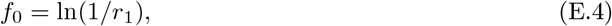

using the bisection method [65].
4. Identify which reaction is going to occur next by picking the reaction index *j* to satisfy the inequality

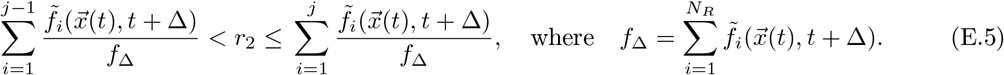
5. According to which reaction has occurred, update the species vector, 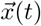.
6. Update the time by replacing *t* with *t* + Δ.
7. If *t* ≤ *t*_max_, then go back to step 2; otherwise, end.

**Figure F.1:**
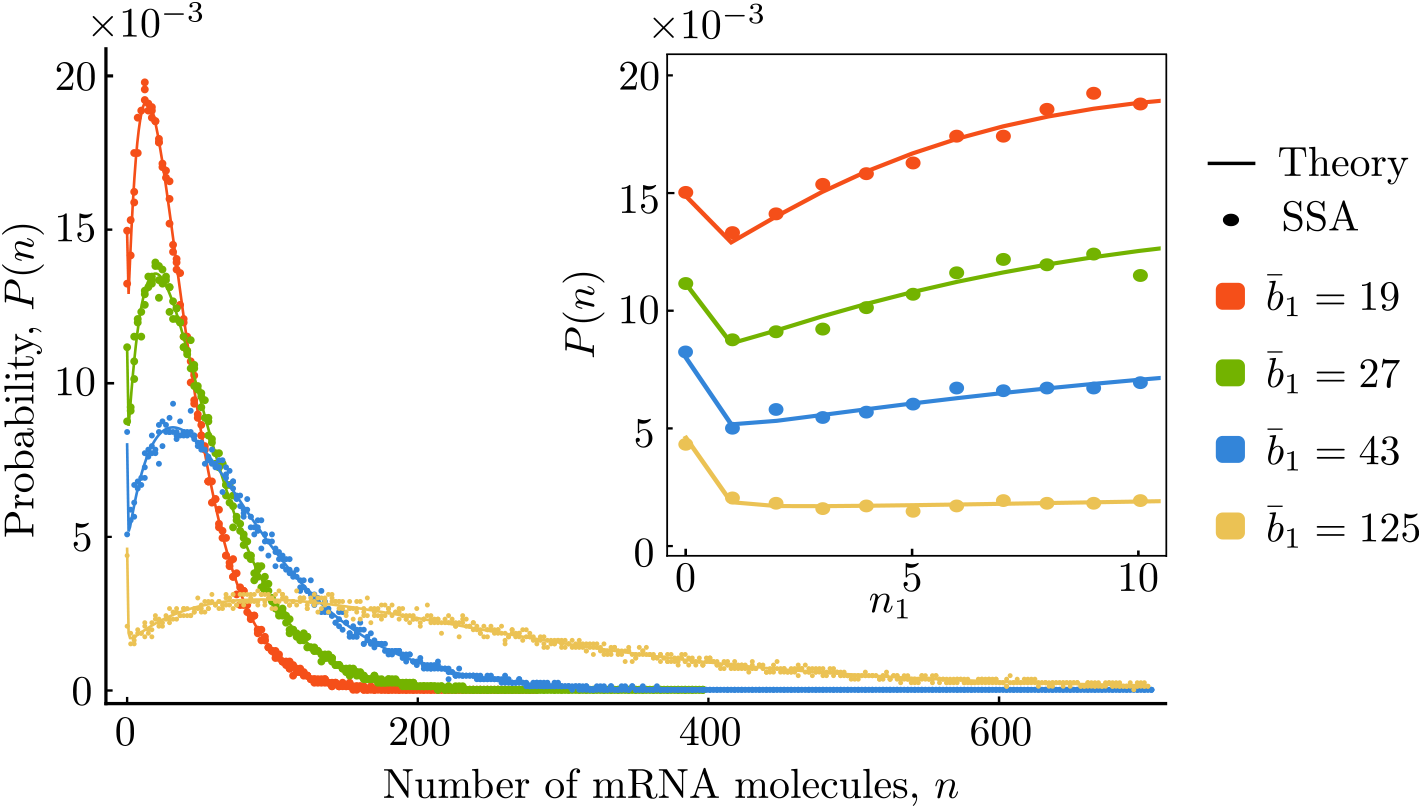
The mRNA distribution from the EM and the RM can display bimodality. We show the mRNA distribution for species *M*_1_ from the EM, as found from stochastic simulation (SSA; points) and predicted exactly by our theory (Eq. (23) together with Eq. (21); solid lines). Note that the above is identical to what is found in the RM, since the two models agree exactly in their prediction of the distribution for *M*_1_. We illustrate results for four different values of the mean burst size, 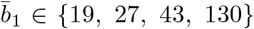. The parameter values that have been used are *ξ* = 0 (cyclo-stationary limit), 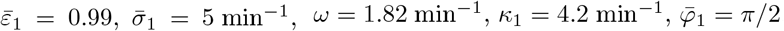, and *t* = 9.2 min. The inset shows a zoomed-in version of the distribution for small numbers of mRNA molecules.

### F Derivation of the exact mRNA distribution for the EM

In this section, we consider the simple reaction scheme given in Eq. (7) for the EM, and we derive its exact time-dependent mRNA distribution. Throughout the following derivations, we consider *j* as some user-input (fixed) parameter which depends on the life-cycle stage of interest. We begin with the PDE for the probability generating function given in Eq. (20), which we convert to the following system of ODEs by using the method of characteristics:

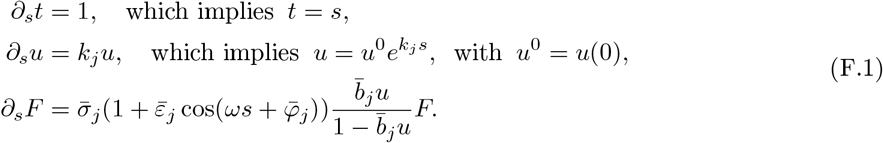

The last ODE in the above system can be rewritten as

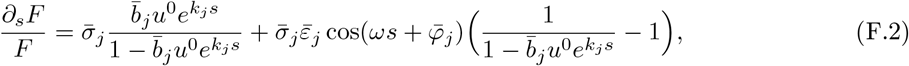

which admits the solution

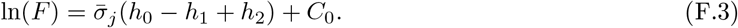

Here, *C*_0_ is some constant of integration to be determined later, with

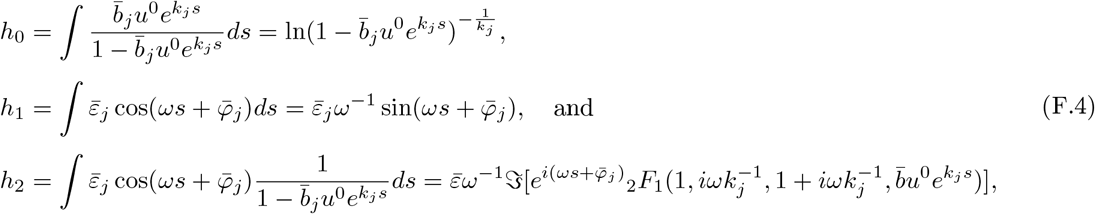

which are well defined when *ω* ≠ 0 and *k*_*j*_ ≠ 0. Here, _2_*F*_1_ is the hypergeometric function of the second kind [53, 54] and ℑ[*z*] denotes the imaginary part of a complex number, *z*. Note that in the expression for *h*_2_, we have used the identity 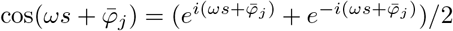, as well as the following identity for hypergeometric functions:

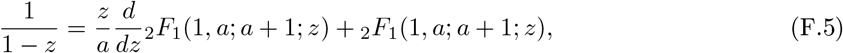

which one can easily derive from Equation 15.5.20 in [53]. Next, we need to determine the constant of integration, *C*_0_. The generating function *F* has to satisfy the initial condition *F*(*s* = 0) = 1 or, equivalently, *F*(*t* = 0) = 1, which stems from the initial condition, *P*(*n* = 0; *t* = 0), as there are zero mRNA molecules in the system at time zero. Applying this initial condition to Eq. (F.3), we have that

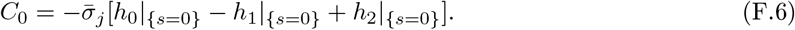

Substituting Eq. (F.6) into Eq. (F.3) and simplifying, we obtain the following solution:

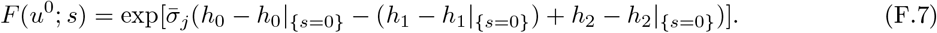

Next, we apply the inverse transformation *s* = *t* and 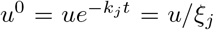 to obtain our final expression for the generating function,

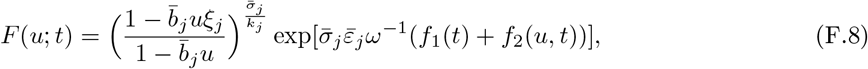

where

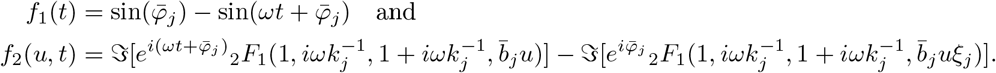

The mRNA distribution is then found from the formula 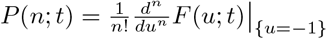.

